# NDR1 and the Arabidopsis Plasma Membrane ATPase AHA5 are Required for Processes that Converge on Drought Tolerance and Immunity

**DOI:** 10.1101/2021.06.10.445978

**Authors:** Yi-Ju Lu, Huan Chen, Alex Corrion, Ilker Buyuk, Pai Li, Saroopa Samaradivakara, Ching Man Wai, Hikaru Sakamoto, Patrícia Santos, Robert VanBuren, Yongsig Kim, Brad Day

## Abstract

NON-RACE-SPECIFIC DISEASE RISISTANCE1 (NDR1) is a key component of plant immune signaling, required for defense against the bacterial pathogen *Pseudomonas syringae*. Plant stress responses have overlapping molecular, physiological, and cell biology signatures, and given the central role of NDR1 during biotic stress perception and signaling, we hypothesized that NDR1 also functions in abiotic stress responses, including in a role that mediates signaling at the plasma membrane (PM) - cell wall (CW) continuum. Here, we demonstrate that NDR1 is required for the induction of drought stress responses in plants, a role that couples stress signaling in an abscisic acid-dependent manner. We show that NDR1 physically associates with the PM-localized H^+^-ATPases AHA1, AHA2, and AHA5 and is required for proper regulation of H^+^-ATPase activity and stomatal guard cell dynamics, providing a mechanistic function of NDR1 during drought responses. In the current study, we demonstrate that NDR1 functions in signaling processes associated with both biotic and abiotic stress response pathways, a function we hypothesize represents NDR1’s role in the maintenance of cellular homeostasis during stress. We propose a role for NDR1 as a core transducer of signaling between cell membrane processes and intercellular stress response activation.

## Introduction

Crop productivity is impacted by a myriad of abiotic and biotic stresses, and among these, water availability and disease pressure are the two primary factors limiting performance and yield^1, 2^. In recent years, many of the genetic and physiological processes that underpin plant resilience in response to both abiotic and biotic stress, as well as the convergence of signaling mediating growth, development, and response to the environment, have been defined^3^. Indeed, work in this area has demonstrated that plants rely on highly conserved genetic mechanisms that underscore growth and development^4^, and environmental stress tolerance^5^ during immune signaling. Particularly noteworthy is the innate ability of plants to respond to, and often overcome, the onset of multiple, simultaneous, stress events^6^. Related to the work described herein, several recent studies have identified substantial signaling overlap between a battery of stress signaling processes, including the convergence of processes governing immunity, adaptation to changes in temperature, and water and nutrient availability^1^.

Among the best-characterized mechanisms of water stress tolerance and adaptation are signaling processes linked to reductions in transpiration, a process mediated by the rapid closure of stomata on the leaf surface via the action of the plant hormone abscisic acid (ABA^7,8^). Additionally, and as a mechanism broadly classified as protection and maintenance of cellular homeostasis, the modulation of membrane dynamics is a primary physiological process associated with dehydration-associated responses in plants^9^. These include, for example, the regulation of signaling associated with the maintenance of cellular membrane homeostasis and metabolic dysfunction^10,11^. As a second, downstream layer of stress response signaling during water deficit, plants also activate numerous dehydration responsive genes^12,13^, activate the production of LEA/dehydrin-type proteins^14^ and various molecular chaperones^15^, as well as relying on cell detoxification mechanisms for the removal of reactive oxygen species (ROS^16^). Not surprisingly, disease resistance signaling also engages similar cellular processes and signaling mechanisms.

Abscisic acid is one of several common denominators linking immunity and abiotic stress signaling^17^. Indeed, several studies in this area have demonstrated that application of exogenous ABA enhances disease susceptibility^18,19,20^, while insensitivity to ABA enhances disease resistance^21,22^. As a regulator of water deficit responses, the accumulation of abscisic acid (ABA) is required for the activation of abiotic stress mechanisms, including downstream gene activation and cellular physiological changes. Among the most important downstream cellular changes in response to rapid accumulation of ABA are rapid turgor presser changes to close stomata. In support of the research described herein, several recent studies have identified significant overlap between water stress response signaling and the regulation of plant defense and immunity^23,24^.

In the current study, we demonstrate that NDR1 (NON RACE-SPECIFIC DISEASE RESISTANCE-1) is required for the robust transcriptional activation of key ABA-associated stress signaling responses in plants, including those associated with both biotic and abiotic signaling mechanisms. Using a combination of physiological, genetic, and transcriptome-based analyses, we further show that a loss of *NDR1* results in a block in stomatal-based mechanisms impacting the response to abiotic and biotic stress signaling. As a mechanism underpinning these responses, we demonstrate that NDR1 associates with the plasma membrane-localized H^+^-ATPases AHA1, AHA2, and AHA5, the function of which is to negatively regulate the activity of the guard cell-localized ATPase. We further demonstrate that overexpression of *NDR1* leads to enhanced tolerance to water deficit, suggesting a role for NDR1 in the regulation of processes mediating cellular response, and tolerance, to drought. In fulfillment of a broader role of NDR1, the data presented herein supports a function for NDR1 as a signaling hub required for response to stress.

## Results

Previous work demonstrated that a loss of *NDR1* and a concomitant increase in electrolyte leakage is correlated with a loss in focal adhesion between the plasma membrane and cell-wall^25^. Not surprisingly, similar focal adhesion mechanisms have been described in response to abiotic stress elicitation^26^. To determine if NDR1 functions in abiotic stress perception and signaling, we first evaluated the *ndr1* mutant for response to water deficit. As shown in Fig. 1a and Supplementary Fig. 1, WT Col-0, the *ndr1* mutant, the *NDR1* native promoter-drive complementation line (*ndr1*/*P_NDR1_*∷*T7-NDR1^25^*), and the *NDR1* overexpression line (*ndr1*/*35S*∷*NDR1^27^*) showed similar responses at 14 days-post water withholding (dpw). However, at 16 dpw, the expression of phenotypes associated with water deficit (e.g., leaf wilting) became pronounced in the *ndr1* mutant. Interestingly, under the same conditions, *NDR1* overexpression plants (i.e., *ndr1*/*35S∷NDR1*) did not show outwardly visible signs of drought stress at 21 dpw (Fig. 1a and Supplementary Fig. 1), suggesting that overexpression of *NDR1* reduces drought-associated phenotypes.

**Fig. 1.**
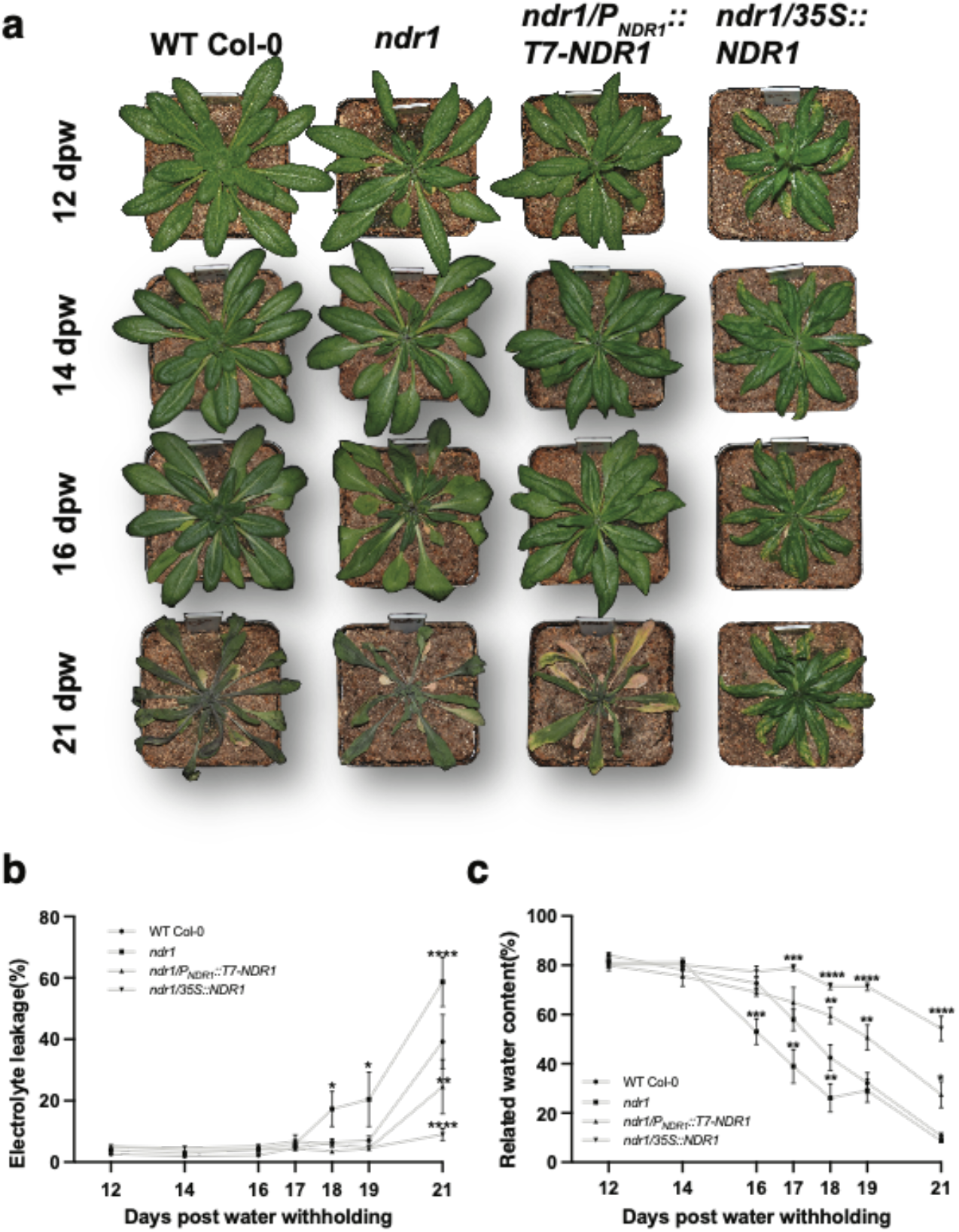
The *ndr1* mutant displays enhanced susceptibility to drought. **a,** Progression of drought response phenotypes from 12 to 21 days post-water withholding (i.e., days post-water-withholding, dpw). Representative images from the full time-course are shown. Additional timepoints are shown in Supplementary Fig. 1. **b,** Quantification of electrolyte leakage from drought exposed plants shown in (a). Electrolyte leakage measurements were collected at each indicated time-point, including those shown in Supplementary Fig. 1. Data points and error bars are the average ± SEM of collections from 5 independent leaves with three independent biological repeats (n ≥ 9). **c,** Quantification of relative water content (RWC) from drought exposed plants shown in **a**. Leaves sampled for determining water loss were collected at each indicated timepoint. Data points and error bars are the average ± SEM of collections from 5 independent leaves with three independent biological repeats (n ≥ 10). Statistical analysis was performed using a two-way ANOVA. Asterisks represent significance differences between different plant genotypes at each day. *P* values ≤ 0.05 were considered significant, where \**P* < 0.05, \*\**P* < 0.01, \*\*\**P* < 0.005, and \*\*\*\**P* < 0.001.

To further evaluate the physiological impact of water deficit and the onset of drought stress across genotypes, we first monitored electrolyte leakage and relative water content (RWC) over the drought timecourse. As shown, from Day 12 until ~16 dpw, all plant genotypes maintained a steady-state level of electrolytes (Fig. 1b) and RWC (Fig.1c). However, from 17-21 dpw, drought phenotypes became more pronounced, along with increased electrolyte leakage and a significant decrease in RWC. For example, at 18 dpw, the *ndr1* mutant showed an approximate 20% increase in electrolyte leakage compared to WT Col-0 and the *NDR1*-expression lines. Consistent with this, at 21 dpw, electrolyte leakage reached ~60% in the *ndr1* mutant, which represents a 20% increase compared to WT Col-0 and a ~50% increase in leakage compared to the *ndr1*/*35S*∷*NDR1* overexpression line (Fig. 1b). Similarly, a decrease in RWC was first observed in the *ndr1* mutant at 16 dpw, and the RWC in WT Col-0 and the *ndr1* mutant dropped ~50% more than in the *NDR1* overexpression line, which maintained RWC of approximately 60% at 21 dpw (Fig. 1c). In total, these data suggest that *NDR1* plays an important role in the response to low water availability and drought stress. As shown in Supplementary Fig. 2, WT Col-0, the *ndr1* mutant, and the *ndr1*/*P_NDR1_*∷*T7-NDR1* complementation line did not recover from the effects of drought (25 dpw) following a 2 day post recovery (soil saturation) period. However, in the *ndr1*/*35S*∷*NDR1* overexpression line, recovery was observed, including ultimate bolting, flowering, and the generation of viable seeds.

To exclude the possibility that T-DNA insertion-site effects underscore the observed response(s) in the *ndr1*/*35S∷NDR1* overexpression line, we assembled a draft genome of the T-DNA mutant to determine the location and copy number of the *NDR1* transgene insertion. As shown in Supplementary Fig. 3, the right border of the *35S∷NDR1* transgene insertion site was located on chromosome 1 at nucleotide (nt) position 5,049,056, and the other end of the plasmid sequence (i.e., left border) was identified at nt 5,049,121. As a result of this insertion event, a 65 nt segment was eliminated. The *35S∷NDR1* transgene lies between two genes: *AT1G14687* and *AT1G14688*. The first gene (*AT1G14687*) encodes a zinc finger homeodomain 14 transcription factor located upstream of the insertion site (Chr1: 5,047,782-5,048,752). The second gene (*AT1G14688*) is located 404 bp downstream of the insertion site (Chromosome 1: nt 5,049,526-nt 5,050,983); this gene encodes an E3 ubiquitin ligase. This analysis supports our assertion that a single insertion event of the *35S∷NDR1* transgene into an intergenic region is causal to the observed drought tolerance phenotype.

The drought-associated phenotype observed in the *ndr1* mutant suggests a likely association with the ABA signaling cascade^28^. To investigate this, and to identify possible miscues in ABA biosynthesis and signaling in the *ndr1* mutant, we first employed an ABA-based seed germination assay, previously demonstrated as a high-throughput marker for monitoring ABA sensitivity^28^. WT Col-0 and *ndr1* mutant seeds had nearly identical germination rates over a 10-day timecourse in the absence of exogenously applied ABA. However, in the presence of 0.5 μM (Fig. 2a), 1 μM (Supplementary Fig. 4a), and 2 μM ABA (Supplementary Fig. 4b), both WT Col-0 and the *ndr1* mutant showed a 1-3 day delay in the germination rate, similar to that previously observed for WT Col-0^29^. However, over the course of the 10 day experiment, the *ndr1* mutant consistently showed statistically significant higher levels of germination in the presence of exogenous ABA, as compared to WT Col-0. The results suggest that NDR1 has a role in ABA-mediated regulation of seed germination.

**Fig. 2.**
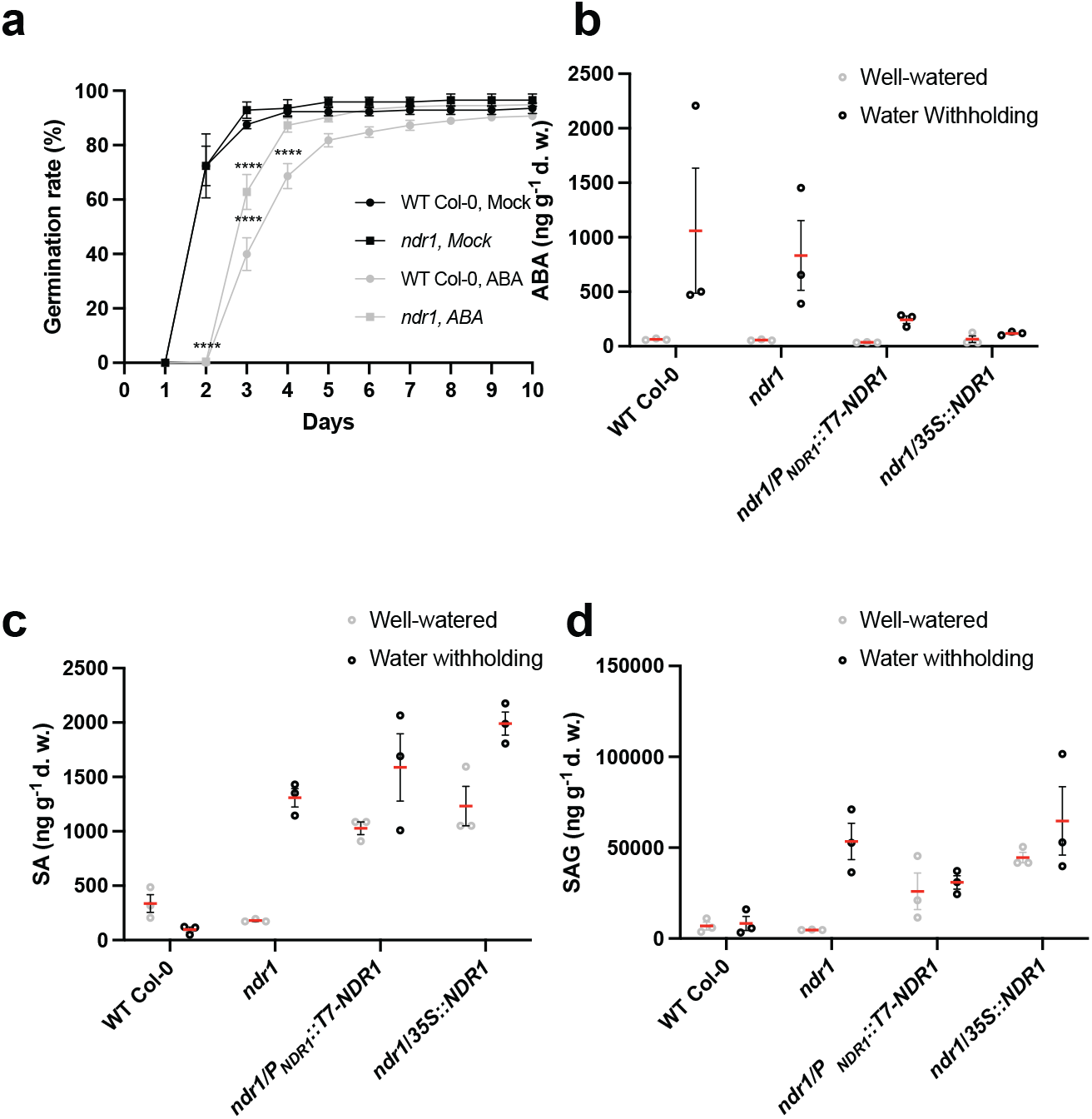
*ndr1* mutant plants are less sensitive to ABA treatment. **a,** Germination assay in the presence of 0.5 μM abscisic acid (ABA), **b,** Quantification of abscisic acid (ABA), **c,** salicylic acid (SA) and **d,** SA-glycosides (SAG) from the leaves of 4-to-5-week-old plants at 16 days post-water withholding (dpw). All data points represent samples collected from 15 individual plants. Statistical analysis was performed using a two-way ANOVA, where *P* ≤ 0.05. Bars indicate standard error, Red lines represent mean values.

To determine if a reduction in root growth is responsible for the observed drought phenotype in the *ndr1* mutant, we next evaluated root phenotypes during exogenous exposure to increasing concentrations of ABA, polyethylene glycol (PEG 6000; i.e., osmotic stress elicitor^30^), high salt (NaCl), and mannitol (i.e., osmotic stress elicitor^31^). As shown in Supplementary Fig. 5, root elongation in the *ndr1* mutant was similar to WT Col-0 in the presence of increasing concentrations of ABA (Supplementary Fig. 5a), up to 20% PEG 6000 (Supplementary Fig. 5b), 150 mM NaCl (Supplementary Fig. 5c), and 300 mM mannitol (Supplementary Fig. 5d) and under well-watered conditions (Supplementary Fig. 5e). Based on these observations, we surmise that root length is not a primary factor underpinning the observed drought sensitivity in the *ndr1* mutant.

To define the relationship between hormone biosynthesis and stress signaling activation in the *ndr1* mutant, we next evaluated the accumulation of key phytohormones (e.g., abscisic acid, ABA; jasmonic acid, JA; salicylic acid, SA) during conditions of water deficit. As shown in Fig. 2b, ABA levels remained the same under well-watered conditions across all genotypes, confirming that the loss of *NDR1* does not affect the biosynthesis of ABA. However, at 16 days of water withholding, both WT Col-0 and *ndr1* mutant plants showed significant increases in the accumulation of ABA compared to well-watered plants. However, the *ndr1*/*P_NDR1_*∷*T7-NDR1* and the *ndr1*/*35S∷NDR1* showed a slightly reduced, albeit significant, increase in ABA. These data suggest NDR1 suppresses ABA biosynthesis in response to water stress. As an additional point of reference, we also observed similar *NDR1* mRNA levels under both control and water withholding which indicates drought stress does not impact *NDR1* gene expression (Supplementary Fig. 6).

As a point of convergence in drought response and plant defense hormone signaling in the *ndr1* mutant, we asked if the accumulation of SA was affected by conditions of limited water availability. Under well-watered conditions, the endogenous levels of SA (Fig. 2c) and the inactive SA-glycosides (SAG) (Fig. 2d) correlated with the relative mRNA accumulation of *NDR1* in each of these lines (Supplementary Fig. 6). SA and SAG levels across all genotypes support our previously published data that the *ndr1* mutant is more susceptible to *Pst* DC3000, a response that is linked to reductions in the defense hormone SA^25^. Additionally, these data suggest that overexpression of *NDR1* leads to a de-repression of SA biosynthesis. Interestingly, under drought stress, both SA and SAG levels significantly increased in the *ndr1* mutant at 16 dpw (Fig. 2 c and d). In the *NDR1* native promoter complementation line, SA levels were modestly enhanced in comparison to WT Col-0 at 16 dpw; a similar trend was observed in the *35S∷NDR1* line, with an approximate 25% increase in SA in response to water withholding (Fig. 2 c and d).

The balance between ABA biosynthesis and catabolism is a critical mechanism that ensures rapid activation, and attenuation, of signaling in response to a battery of external environmental stressors^32^. Based on the data presented in Fig. 2, we hypothesized that the observed phenotype(s) in the *ndr1* mutant under water deficit are likely not due to a block in the accumulation of key defense and abiotic stress signaling hormone. However, we cannot exclude the possibility that the induction of key ABA biosynthesis genes, as well as ABA responsive marker genes, are not affected, and thus causal to the observed phenotypes. To investigate this, we evaluated the mRNA accumulation of a suite of genes associated ABA biosynthesis and metabolism using real-time qPCR. As shown in Supplementary Fig. 7a, mRNA accumulation of the upstream enzyme in ABA biosynthesis, *ZEAXANTHIN EPOXIDASE* (*ABA1*), is elevated to a similar level during drought conditions compared to well-watered controls in WT Col-0, the *ndr1* mutant, the *ndr1*/*P_NDR1_*∷*T7-NDR1*, and the *ndr1/35S∷NDR1* lines. We also examined additional downstream enzymes that are important for ABA biogenesis. For example, a significant increase in *9-CIS-EPOXYCAROTENOID DIOXYGENASE* (*NCED3*) is detected in WT Col-0 and the *ndr1* mutant under drought stress (Supplementary Fig. 7b). The accumulation of *XANTHOXIN DEHYDROGENASE* (*ABA2*), which encodes a key enzyme in the early conversion steps of xanthoxin to ABA aldehyde, remained to the similar levels and was significantly reduced under water withholding in the *ndr1* mutant (Supplementary Fig. 7c). The expression level of *ABA2* implies a positive correlation with *NDR1* mRNA levels in all tested lines (Supplementary Fig. 6). Interestingly, we found the mRNA expression of ABA biosynthesis aldehyde oxidase (*AAO3*) is significantly induced in WT Col-0 and the *ndr1* mutant but not the *NDR1* over-expressor lines (Supplementary Fig. 7d). Moreover, similar mRNA expression patterns were also observed when we measured *RD29B* and *EARLY RESPONSIVE TO DEHYDRATION* (*ERD4*;^33^), both of which are responsive to desiccation and dehydration (Supplementary Fig. 7e, f).

As shown in Fig. 1, the *ndr1* mutant exhibited a striking decline in RWC in response to water deficit at 16 dpw. Coincident with this, we detected a slight, yet significant, decrease in *ABA1* mRNA in the *ndr1* mutant compared to WT Col-0. Conversely, by day 21, and in conjunction with the onset of severe drought-associated phenotypes (e.g., Fig. 1), we observed a significant increase in the mRNA accumulation of *ABA1* in the *ndr1* mutant. At the same time, mRNA accumulation of *ABA2* shifted from being significantly downregulated at 16 dpw in the *ndr1* mutant (*P* < 0.05). Taken together, and in agreement with our quantitative evaluation of RWC over the timecourse of water withholding (Fig. 1c), we propose that 16 dpw represents a tipping point for the *ndr1* mutant and WT Col-0 plants, respectively.

Drought response signaling in plants is regulated by both ABA-dependent and ABA-independent pathways, both of which share convergent regulatory mechanisms^34^. For example, many drought-associated genes are regulated by rhythmic changes in the circadian clock, a genetic network that drives plant develop, response to pathogens, and control of hormone biosynthesis^34,35^. Stomata are critical components of water deficit signaling, functioning in large part by regulating the diurnal rate of transpiration. As a point of entry for bacterial pathogens, stomata aperture is directly correlated with infection and pathogen colonization^36^. To determine if a loss of *NDR1* results in alterations in the clock, we evaluated the diurnal gating of stomatal guard cells as a tractable, predictable, output of clock function, and moreover, diurnal hormone oscillations in hormone accumulation. As shown in Supplementary Fig. 8a, we observed that the *ndr1* mutant had increased stomatal aperture, compared to WT Col-0, at Zeitgeber time 5 h and 7 h; as indicated, this period corresponds to daylight hours. During Zeitgeber time 15 h – 23 h, the *ndr1* mutant displayed an increase in guard cell aperture (i.e., increased opening) when compared to WT Col-0. As predicted, overexpression of *NDR1* resulted in enhanced guard cell closure over the duration of the 24 h diurnal cycle. In agreement with this, the mRNA expression of key circadian marker genes was also affected in the *ndr1* mutant at timepoints coincident with the hours preceding daylight. Further, the accumulation profiles of ABA and SA over the 24 h diurnal cycle were the same across all 3 genotypes (Supplementary Fig. 9).

Based on our observations detailed above, we next asked if NDR1 is required for the convergence of signaling between pathogen perception and stomata response. Leaf punches from WT Col-0 and the *ndr1* mutant were treated with 10 μM ABA. As shown in Fig. 3a, at 1 h post-treatment, WT Col-0 stomata exhibited a significant (*P* < 0.0001) reduction in aperture width following ABA treatment, while *ndr1* did not show a change in aperture as compared to mock-treated controls. Similar to the absence of stomatal closure in response to ABA treatment, we also observed that exogenous application of polyethylene glycol (PEG) (Fig. 3b), mannitol (Fig. 3c), and hydrogen peroxide (H_2_O_2_) (Fig. 3d) did not induce stomatal closure in the *ndr1* mutant.

**Fig. 3.**
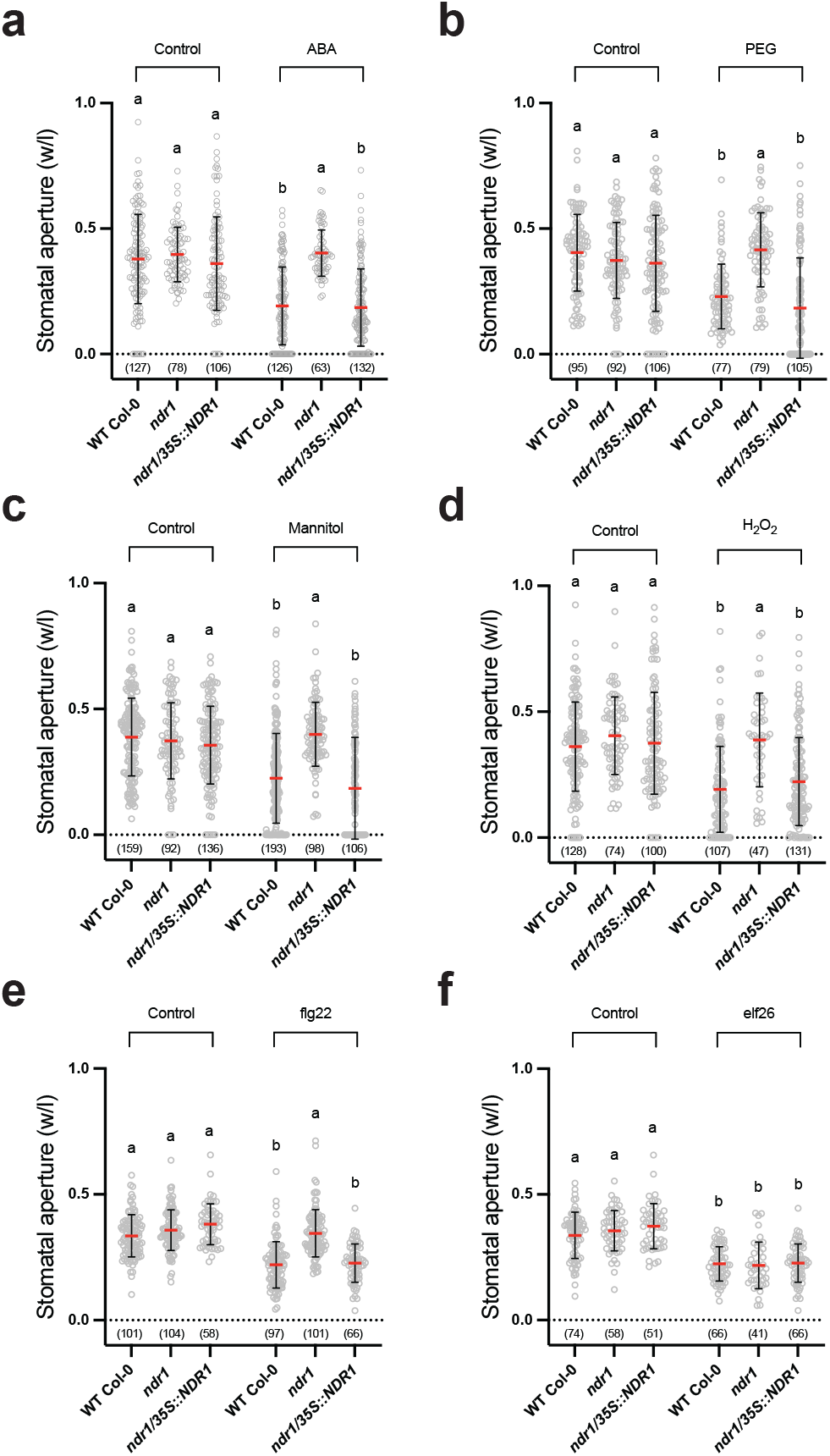
Stomatal closure in the *ndr1* mutant is impaired, with differential responses to ABA and PAMPs. Four-to-five-week old plants were subjected to either abiotic or biotic stress elicitation for one hour before imaging and guard cell aperture evaluation. As shown, *ndr1* guard cells do not respond to **a,** 10 μM ABA, **b,** 20% PEG, **c,** 300 mM mannitol, **d,** H_2_O_2_, **e,** 100 nM flg22, **f,** 100 nM elf26. The *ndr1* guard cells do respond to **d,** elf26. Statistical analysis was performed using a two-way ANOVA, where *P* ≤ 0.05. Bars indicate standard error, Red lines represent mean values.

Immune signaling downstream of bacterial flagellin (i.e., flg22) perception is compromised in the *ndr1* mutant^25,37^. To investigate the implication of reduced flg22 signaling and/or general PAMP (pathogen-associated molecular pattern) perception in the *ndr1* mutant, we evaluated stomatal aperture response following flg22 treatment. Whole leaf punches from WT Col-0 exposed to flg22 showed rapid guard cell closure (ca. 1 h; Fig. 3e), whereas guard cells from the *ndr1* mutant remained open following flg22 treatment. To determine if additional PAMP treatment and perception responses were affected in the *ndr1* mutant, we also evaluated guard cell response following elf26 elicitation^38^. Surprisingly, in response to elf26 treatment, stomatal closure in the *ndr1* mutant was similar to that observed in WT Col-0 (Fig. 3f).

Leaf water content is a major contributor to the restriction of *Pst* DC3000 growth during gene-for-gene mediated resistance in *Arabidopsis* due to the coupling of restricted vascular activity and water loss through the stomata at the site of infection^39^. Indeed, previous work showed that *Pst* DC3000 induces significantly lower water potentials during incompatible interactions than during compatible interactions^40^. Based on these observations, we hypothesized that the *ndr1* mutant would be more susceptible to bacterial infection under conditions of drought stress. To test this, we evaluated the impact of drought on the development of disease and enhanced bacterial growth following pathogen infection of 4-week-old WT Col-0 and *ndr1* mutant plants subjected to water withholding. As expected, the difference in *Pst* DC3000 growth in *ndr1* mutant plants under well-watered conditions was approximately 1.5 log CFU cm^−1^ higher than in WT Col-0 (Supplementary Fig. 10a). Under conditions of water-withholding, a similar differential (ca. 1.5 log CFU cm^−1^) in *Pst* DC3000 growth was observed in the *ndr1* mutant as compared to WT Col-0. In agreement with the role of NDR1 as a positive regulator of plant defense, the *in planta* growth of *Pst* DC3000 in the *NDR1* native promoter-driven and overexpression lines (Supplementary Fig. 6) were similar to that observed in WT Col-0. Interestingly, under drought conditions, plants challenged with the avirulent pathogen, *Pst* DC3000-AvrRpt2, showed a reponse consistent with the hypothesis presented by Wright and Beatie^40^. Indeed, we observed resistance to *Pst* DC3000-AvrRpt2 in WT Col-0 and susceptibility in the *ndr1* mutant under well-watered conditions, as previously described^25,27,41^ (Supplementary Fig. 10b). However, under conditions of water-withholding, growth of *Pst* DC3000-AvrRpt2 in the *ndr1* mutant was ~1 log greater than under well-watered conditions. These data are in agreement with previous studies demonstrating that drought stress impacts bacterial growth during incompatible interactions but not during compatible interactions^39^.

Based on the data presented above, we next asked if the requirement for NDR1 is associated with transcriptional activities of immunity and/or ABA-associated signaling processes. To test this, we conducted a comprehensive mRNA-seq analysis to evaluate gene expression changes in WT Col-0, the *ndr1* mutant, and the *NDR1* overexpression line (*ndr1/35S∷NDR1*) over the timecourse of 0-21 day post water-withholding. Control samples (i.e., well-watered) were collected from all genotypes and at all time points. In total, these datasets were comprised of 129 RNA-seq libraries. To explore gene expression difference between treatment groups and WT Col-0 group, samples were grouped using principle component analysis (PCA), the output of which revealed that gene expression in different groups were primarily separated from each other by time, with distinct grouping at Day 0,12,14,16,17,18, 19, and 21 under water withholding treatments, and at Day 0, 12, 16, and 21 under well-watered treatment (Supplementary Fig. 11).

To establish a baseline of *NDR1*-dependent transcriptional responses (i.e., differentially expressed genes; DEGs (P value < 0.01, |log_2_-fold change| ≥ 1)) across genotypes, we first evaluated gene expression profiles from each of the 3 genotypes from 0-21 day post water-withholding. As a control for stress-responsive DEGs, we also included parallel samples (i.e., each genotype at each timepoint) from well-watered plants. As shown in Figure 4, we first identified mRNAs that were differentially expressed compared to well-watered plants in each group over the drought timecourse in *ndr1* and *ndr1/35S∷NDR1* compared to WT Col-0 plants. We identified significantly more differentially regulated genes in the *NDR1* overexpression line compared to the *ndr1* mutant (Figure 4a; Supplemental Data Set 1 and 2). We also observed the same trend as above in the well-watered treatment (Figure 4b, Supplemental Data Set 3 and 4). Next, we examined the pattern of DEGs among each time points to evaluate the temporal pattern of expression of key genes as a function of genotype and treatment. As shown in Figure 4c, DEG patterns in the *ndr1/35S∷NDR1* overexpression line vs WT Col-0 were similar across the entire drought timecourse, while the pattern varied in the combination of *ndr1* mutant vs WT Col-0. Further analysis of the day-to-day comparison (with Day 0 as a baseline) of DEGs in WT Col-0, the *ndr1* mutant, and *ndr1/35S∷NDR1* overexpression line revealed more DEGs in WT Col-0 than in the *ndr1* mutant and the *ndr1/35S∷NDR1* overexpression line under both water-withholding and well-watered conditions (Supplemental Fig 12a, b). The composition of DEGs among the different plant genotypes was found to be most striking at 12 dpw. Indeed, as the pattern of DEGs in *ndr1* mutant was different from WT Col-0, the *ndr1/35S∷NDR1* overexpression line was able to restore a complementary set of genes, as compared to WT Col-0 (Supplemental Fig 12c, d Supplemental Data Set 5, 6, 7, 8, 9, and 10).

**Fig. 4.**
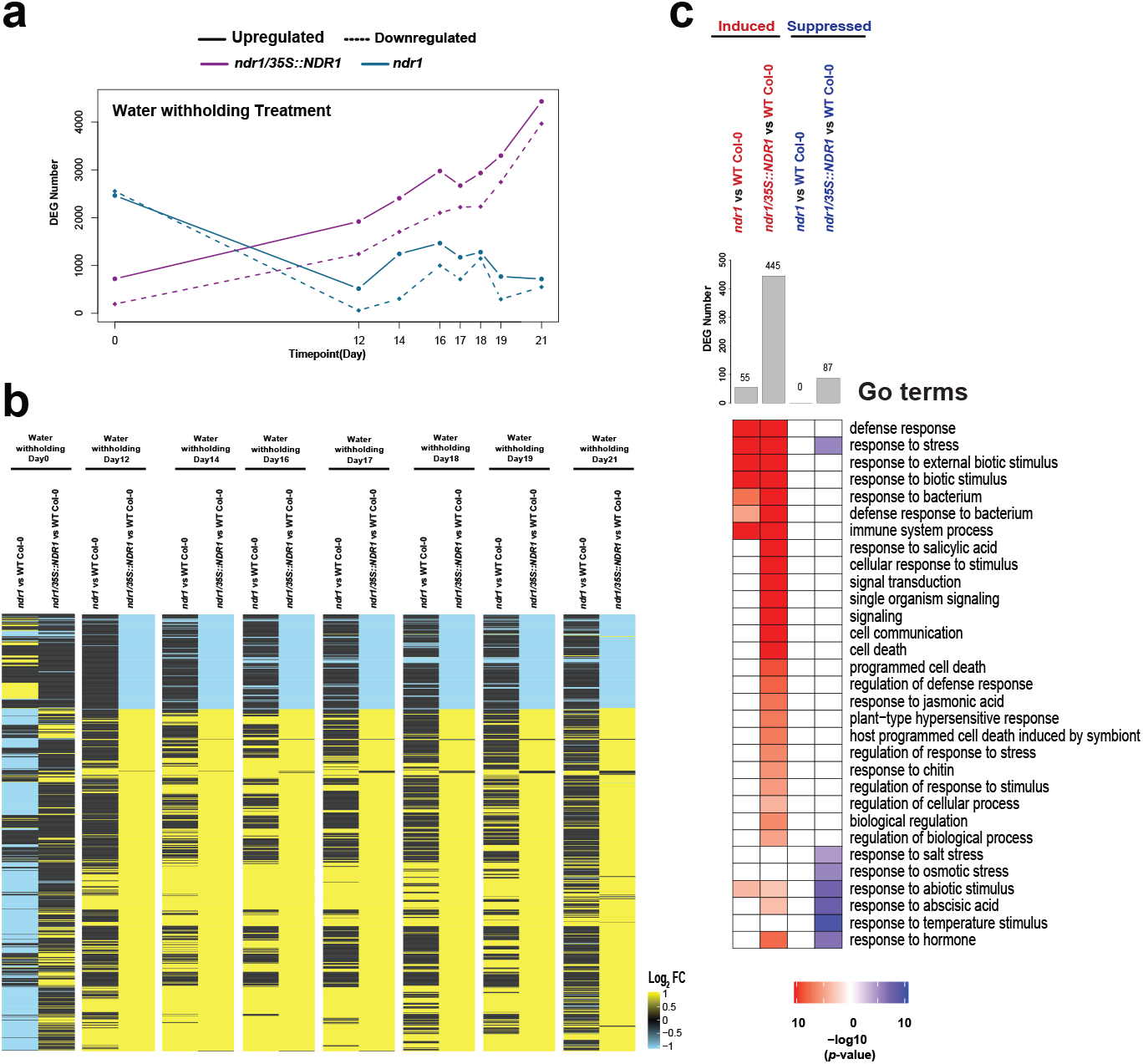
Patterns of differential gene expression and the biological process pathways that are regulated by these DEGs during drought stress. **a,** Expression profile of DEGs under water withholding over the 12-21 day timecourse. **b,** Expression patterns of genes under water withholding condition. **c,** GO terms of differential DEGs between *ndr1* vs WT Col-0 and *ndr1/35S∷NDR1* vs WT Col-0.

To further investigate this, we next identified the DEGs that were constantly up- or downregulated from 12-21 dpw among WT Col-0, *ndr1*, and *ndr1/35S∷NDR1* genotypes. From this, we identified 153 genes that are constantly upregulated in the *ndr1* mutant, and 1319 that are constantly upregulated in the 35S-over-expressor line over the drought timecourse. Additionally, 62 transcripts were identified as constantly downregulated in the *ndr1* mutant, and 831 were constantly downregulated in the 35S over-expressor line (Supplementary Data Set 11, 12, 13, and 14). Among these, 96 transcription factors (TF) belonging to the WRKY, ERF, AP2, and MYB families were identified as either constantly upregulated or downregulated under water deficit and well-watered conditions (Supplemental Fig. 13; Supplemental Data Set 15). Not surprisingly, and in further support of our hypothesis, we found that *WRKY47*, a drought-responsive TF, was also among the list of the above DEGs. Additionally, the mRNA expression levels of *WRKY40* and *WRKY70* were also altered significantly under water withholding conditions (Supplemental Fig. 13).

A gene ontology (GO) enrichment analysis for the 995 DEGs that were constantly upregulated in the *ndr1/35S∷NDR1* line identified GO terms that have been reported to regulate reactions to drought stress and plant immunity including genes response to hormone (i.e., ABA, JA, and SA), stress, intrinsic components of the plasma membrane, and immune system process (Fig. 4c). Among these, we identified 324 downregulated genes in GO term response to osmotic and drought stresses in *ndr1/35S∷NDR1* under water withholding conditions (Fig. 4c; Supplemental Data Set 16 and 17). In support of our assertion that overexpression of NDR1 is responsible for these changes in gene expression profiles, we did not observe similar annotated pathways among the 145 up- and 8 downregulated DEGs from the *ndr1* mutant.

As a final step in our transcriptome analysis, and to associate mRNA expression changes with the known biological activities of the NDR1-dependent immune and abiotic process, we next evaluated the biological processes within each group from 12-21 dpw compared to 0 dpw. Interestingly, we found that changes gene expression occur earlier in *ndr1/35S∷NDR1* than in *ndr1* mutant in response to water withholding treatment (Figure 4c; Supplemental Data Set 18 and 19). This is exciting, as it illustrates that the strongest influence on differential gene expression of water-response processes is linked to the overexpression of *NDR1*. Taken together, these data provide compelling evidence that overexpression *NDR1* plays an important role in the regulation of drought stress-responsive genes under conditions of limited water availability.

Lastly to investigate the relationship between NDR1 function and response to drought as a requirement of ABA signaling, we explored the common and unique DEG in *ndr1* and *ndr1/35S∷NDR1* under water deficit and well-watered conditions, respectively (Supplementary Fig 14a, b). In addition, we cataloged all GO terms associated with our candidate list of DEGs in the above. We found that unique DEGs in *ndr1/35S∷NDR1* under water with-holding treatment were significantly enriched in pathways responding to water and SA. However, DEGs in *ndr1* mutant under water withholding treatment are not enriched in the above biological processes (Supplemental Fig. 14c; Supplemental Data Set 20). Taken together, these data support the hypothesis that immunity and drought process signaling converge as a function of *NDR1* overexpression.

To gain insights into global gene coexpression profiles under water with-holding environments, we performed coexpression analysis in each of the genotypes as a function of water status. In short, this allowed us to categorize genes into coexpression modules and identify the expression levels of module eigengenes (MEs). Using this approach, we identified 15 coexpression modules among WT Col-0, *ndr1*, and the *ndr1/35S∷NDR1* overexpression line at all timepoints (Supplemental Fig. 15; Supplemental Data Set 21-23). As shown in Figure 5a, Module 2 contains genes that are associated with processes associated with plant response to water deficit. In this, we found *LEA14* (At1g01470) and *ERD10* (At1g20450), with a reduced eigengene value in the *ndr1/35S∷NDR1* overexpression line than in WT Col-0 or the *ndr1* mutant. Interestingly, coexpression profiles from Module 3 also highlight a pattern revealing higher expression in defense-related genes in the *ndr1/35S∷NDR1* line, including known stress regulators such as *WRKY70* (At3g45640), *PAD4* (At3g52430), *NPR4* (At4g19660), and *FLS2* (At5g46330). In addition, RNA-seq analysis provided evidence that the expression of *AHA2* was upregulated in *ndr1/35S∷NDR1* plant compared to WT Col-0 across all time points (Figure 5b). These results indicate that NDR1 likely functions in signaling processes required for guard cell gating, a mechanism known to control both biotic and abiotic stress signaling responses.

**Fig. 5.**
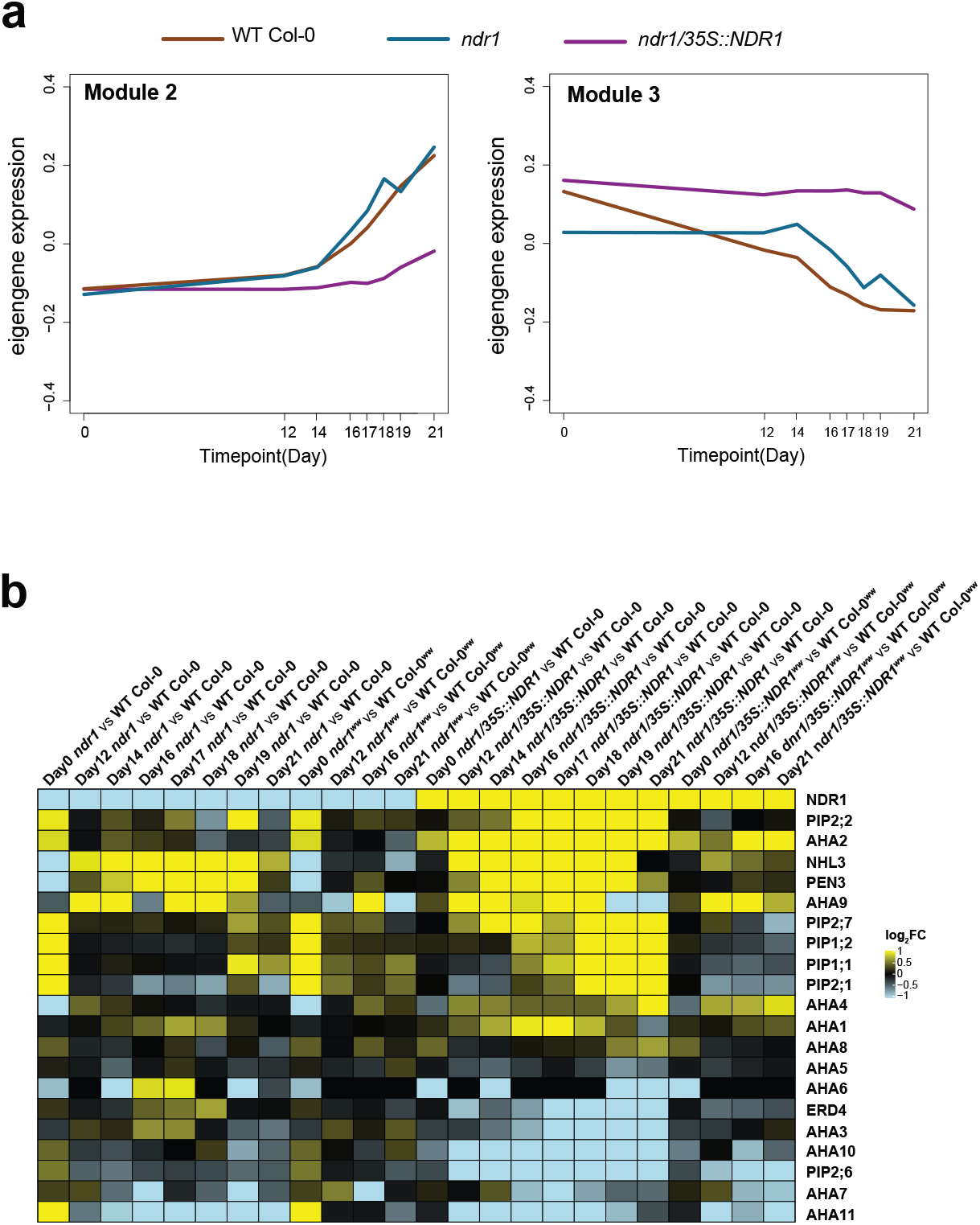
Gene coexpression modules and expression patterns of drought-associated genes during water-withholding. **a,** Eigengene expression modules of WT Col-0, *ndr1*, and *ndr1/35S∷NDR1.* **b,** mRNA accumulation of the mRNAs whose products were identified as NDR1-interacting candidates..

As shown in Figure 5b, we observed an increased in the mRNA accumulation of several membrane-associated H^+^-ATPases, including *AHA2*, *AHA9*, and *AHA4*. Of particular note, and in agreement with the MS/MS data, is the identification of an interaction between NDR1 and membrane H^+^-ATPase, AHA1 (*At2g19860*), which in addition to its role in signaling during abiotic stress response, is required for immune signaling following bacterial pathogen infection^42^. To confirm the specificity of the NDR1-AHA1 interaction, we first performed *in planta* co-immunoprecipitation (co-IP) assays using a transient *Agrobacterium tumefaciens*-*Nicotiana benthamiana* heterologous expression system. To do this, we evaluated the pairwise interactions between Flag-NDR1 and HA-epitope-tagged constructs of AHA1, AHA2, and AHA5. As shown in Figure 6b, we identified in planta interactions between Flag-NDR1 and all three tested AHA proteins, which supports the NDR1-AHA1 IP-MS/MS interaction data shown in Table 1. To further evaluate the interactions between NDR1 and each of the identified AHAs, as well as to provide insight into the cellular address of these interactions, we next used bimolecular fluorescence complementation (BiFC) assays, again confirming the interaction between NDR1 and all tested AHA proteins (Supplementary Fig 16).

**Fig. 6.**
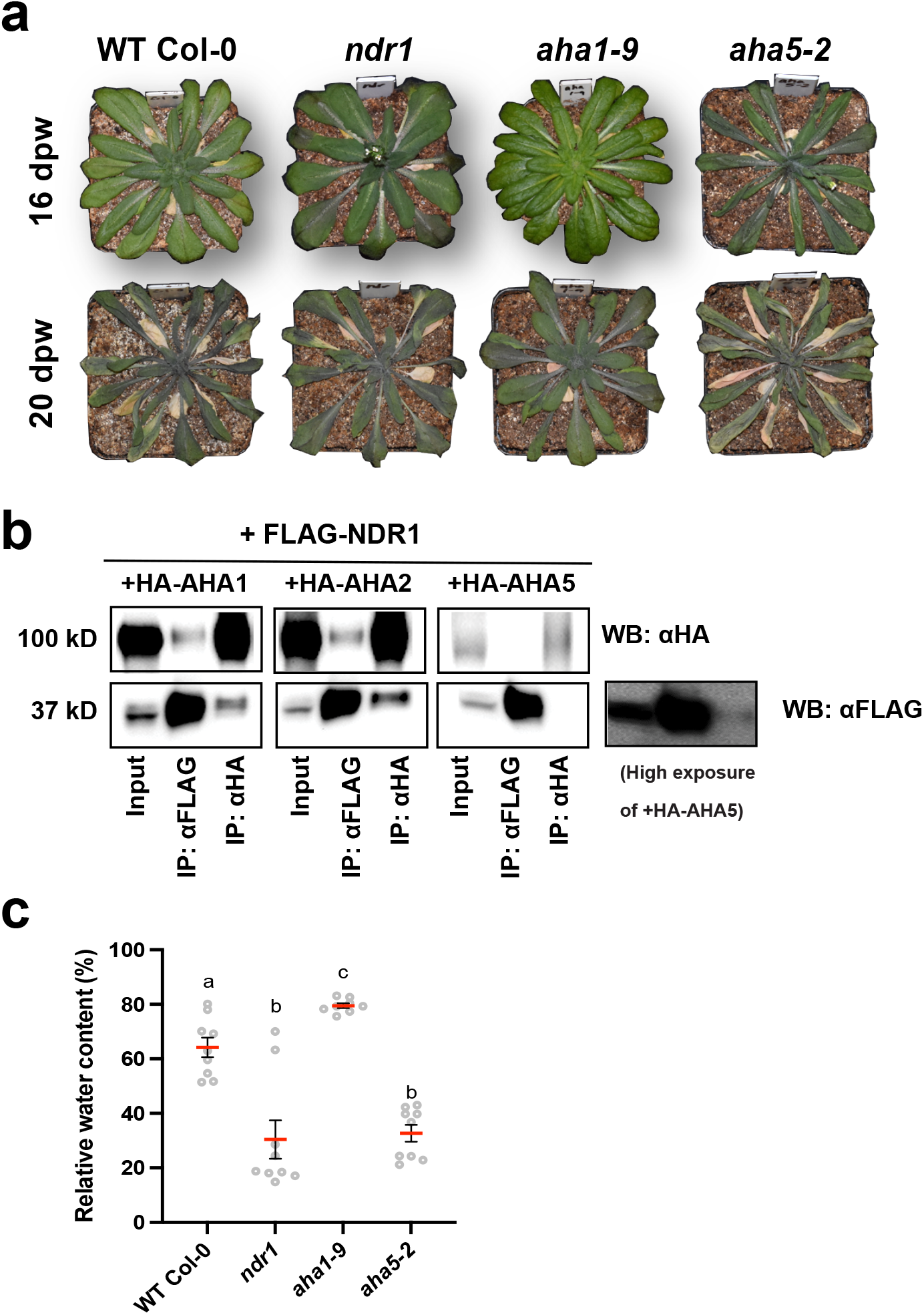
The *aha1-9* mutant shows increased tolerance to water-withholding conditions compared to WT Col-0 and the *ndr1* and *aha5-2* mutants. **a,** Representative images of *Arabidopsis* plants undergoing drought stress over a timecourse of 14-to-20-days post-water withholding. **b,** Co-immunoprecipitation of Flag-NDR1 with HA-AHA1, HA-AHA2, and HA-AHA5 **c,** Relative water content of plants during drought stress at 15 dpw (days post-water-withholding). Statistical analysis was performed using a one-way ANOVA. *P* values ≤ 0.05 were considered significant. Bars indicate standard error, Red lines represent mean values.

**Table 1.** NDR1 interacting proteins identified by immunoprecipitation tandem mass spectrometry. Peptide counts are shown for each of 2 independent biological repeats (i.e., BR-1, BR-2). Protein identification required *P* < 0.05 (MOWSE algorithm), minimum of # peptides. None of the proteins listed were identified in the negative control (i.e., WT Col-0) IP-MS/MS experiment.

| Gene Name | Gene ID | GO Term | Peptide Count |  |
| --- | --- | --- | --- | --- |
|  |  |  | BR-1 | BR-2 |
| NDR1 (control) | AT3G20600 | Non-Race Disease Resistance-1 | 4 | 7 |
| AHA1 | AT2G18960 | H(+)-ATPase-1 | 16 | 15 |
| ERD4 | AT1G30360 | Early-responsive to dehydration stress protein | 12 | 13 |
| PIP2:7 | AT4G35100 | Plasma membrane intrinsic protein | 3 | 7 |
| HIR2 | AT3G01290 | PHB domain-containing membrane-associated protein family | 8 | 3 |
| Remorin1 | AT3G61260 | Remorin family protein | - | 4 |
| PIP1:2 | AT2G45960 | Plasma membrane intrinsic protein | 4 | 4 |
| PIP2:6 | AT2G39010 | Plasma membrane intrinsic protein | 5 | 3 |
| NHL3 | AT5G06320 | NDR1/HIN-like 3 | 2 | 3 |
| HIR1 | AT1G69840 | PHB domain-containing membrane-associated protein family | - | 2 |
| PIP2:2 | AT2G37170 | Plasma membrane intrinsic protein | - | 2 |
| PIP2:1 | AT3G53420 | Plasma membrane intrinsic protein | 4 | 2 |
| PEN3 | AT1G59870 | Plant PDR ABC-type transporter family protein | 2 | 2 |
| PIP1:1 | AT3G61430 | Plasma membrane intrinsic protein | 3 | 2 |

Data presented herein support the hypothesis that NDR1 functions in stomatal osmoregulation immunity through its interaction with plasma membrane-associated H^+^-ATPases. To further test this, we evaluated available SALK T-DNA mutant alleles for response to drought to determine if *AHA1* and *AHA5* phenocopy the *ndr1* mutant (Supplementary Fig. 17). As shown in Figure 6a, we observed that the *aha1-9* mutant was in fact more tolerant to the effects of water withholding, as compared to WT Col-0, whereas *aha5-2* was more susceptible – similar to the *ndr1* mutant. These observations were further supported by a quantitative analysis of the RWC over a short time-course of drought. Indeed, and in agreement with the NDR1-AHA5 protein-protein interaction analysis (Fig. 6b), we observed that the *aha5-2* mutant had similar levels of RWC as the *ndr1* mutant over the timecourse of water withholding, while the *aha1-9* mutant had RWC levels similar to the *NDR1* overexpression line (Fig. 6c).

As shown above, NDR1 is essential for stomatal closure in response to a variety of abiotic stressors, including the differential response to at least 2 well-characterized PAMPs. Based on the identification of a physical interaction between NDR1 and AHAs, we next asked if the loss of function of H^+^-ATPases phenocopies *ndr1* stomatal response following PAMP perception. As shown in Fig. 7, we observed differential responses in the *aha1-9* and *aha5-2* mutant lines. Stomata from WT Col-0 and the *aha1-9* mutant rapidly closed following treatment with ABA (Fig. 7a). However, following ABA treatment, stomata from the *ndr1* and *aha5-2* mutant did not close. A similar result was observed when plants were treated with exogenous flg22 peptide (Fig. 7b). Conversely, plants treated with the elf26 peptide showed rapid guard cell closure (ca. <1 h; Fig. 7c). These data are consistent with the observation that *ndr1* mutant stomata also do not respond to treatment with ABA or flg22 yet does respond to elf26 (Fig. 3e, f).

**Fig. 7.**
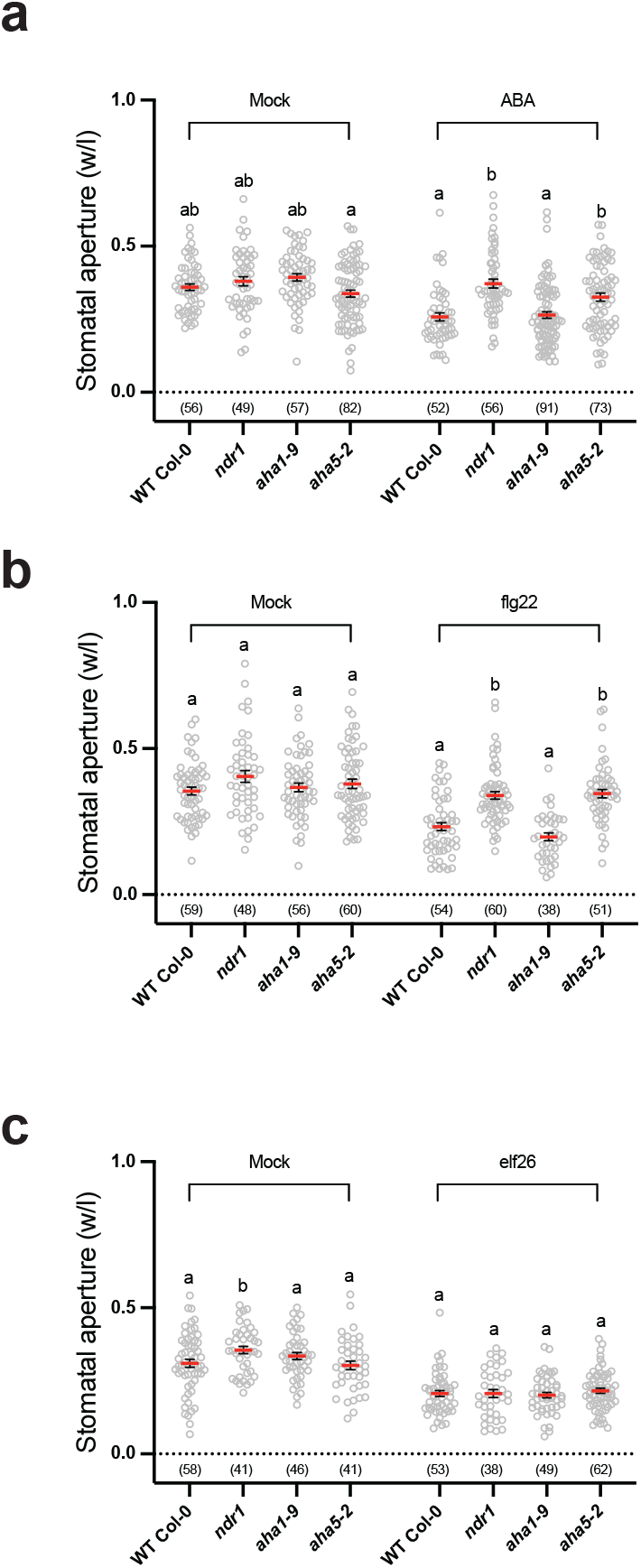
The *aha5-2* mutant displays impaired stomatal closure in response to both abiotic and biotic stress elicitors. Knockout mutants *aha5-2* phenocopies *ndr1* in stomatal response to **a,** ABA, **b,** flg22, and **c,** elf26 treatment. Leaf discs were subjected to stress for 1 hour prior to imagining. Each data point represents 3 biological replicates; > 50 stomata were imaged for each replicate. Statistical analysis was performed using a one-way ANOVA. *P* values ≤ 0.05 were considered significant. Bars indicate standard error, Red lines represent mean values.

As a mechanism controlling stomatal gating in response to phytopathogen infection, it was recently shown that RIN4 acts as a positive regulator of PM H^+^-ATPase activity^42^. Based on our observation that exogenous application of ABA and flg22 are unable to induce stomata closure in the *ndr1* mutant, as well as NDR1’s association with RIN4^43^, we hypothesized that the PM H^+^-ATPase activity in *ndr1* stomata may be affected. To test this, we next monitored PM H^+^-ATPase activity in WT Col-0, the *ndr1* mutant, and the *ndr1/35S∷NDR1* overexpression line, to evaluate NDR1’s role in regulating H^+^-ATPase activity. In support of our hypothesis, we observed an enhancement in the activity of the H^+^-ATPase in the *ndr1* mutant (Fig. 8a), suggesting a potential role for NDR1 in negatively regulating ATPase activity. Our results revealed an increase in H^+^-ATPase activity in the *aha1-9* and *aha5-2* mutants (Fig. 8b). Taken together with the genetic and protein interaction data shown above, we assert that NDR1 functions in guard cell gating through its association with stomatal-localized H^+^-ATPases.

**Fig. 8.**
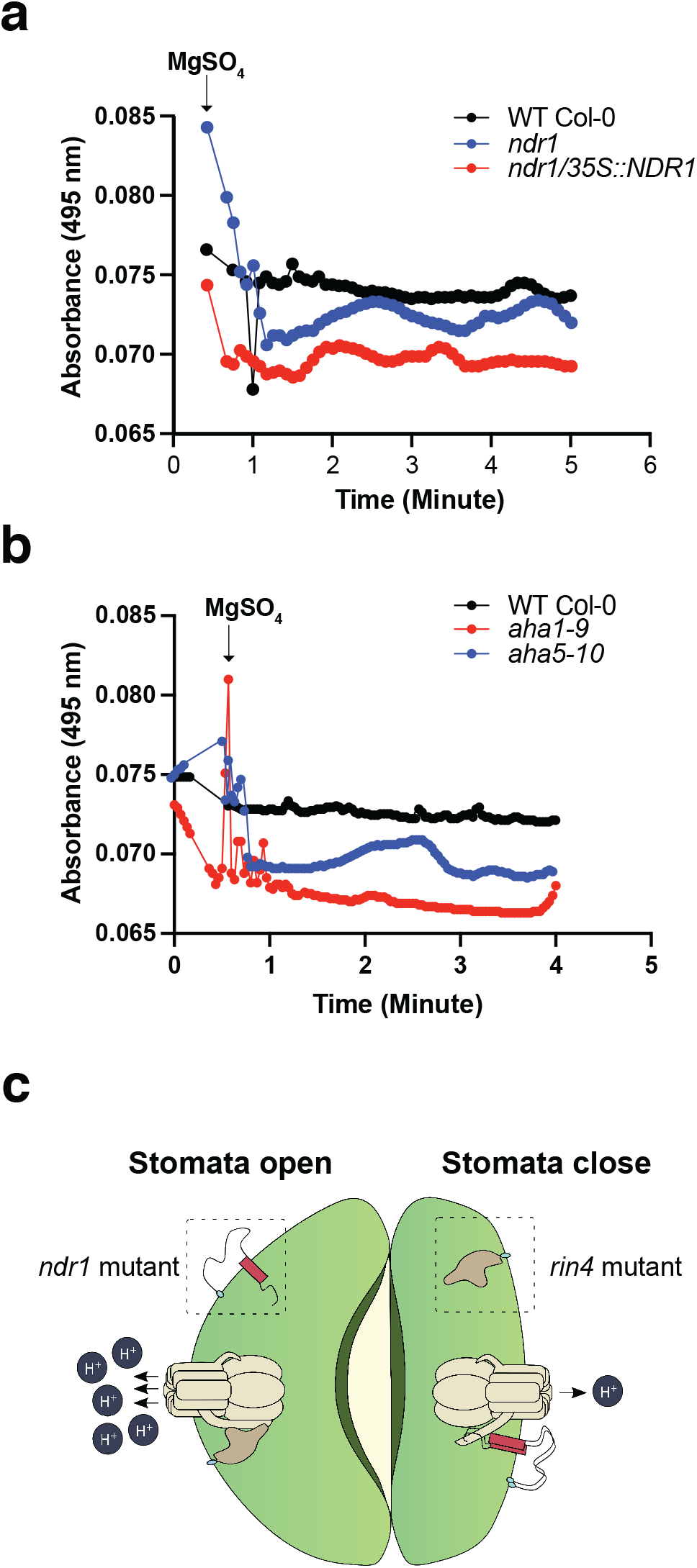
H^+^-ATPase activity is enhanced in the *ndr1* and the *aha1-9* H^+^-ATPase mutant. **a,** WT Col-0 *ndr1* mutant and *35S∷NDR1*, **b,** WT Col-0, *aha1-9*, and *aha5-2* mutant leaf enriched plasma membrane fractions were purified using an aqueous polymer two-phase method, and the assay was conducted on inside-out vesicles. Plasma membrane H^+^-ATPase activity was measured using an acridine orange efflux assay. Absorbance was measured at 495 nm. Experiments were repeated 2-3 times with independent plasma membrane enriched fractions for each replicate. **c,** Model illustrating the link between NDR1, RIN4, and the plasma membrane H^+^-ATPase. NDR1 and RIN4 have opposing functions in both biotic and abiotic stress signaling.

## Discussion

NDR1 is an essential component of the *Arabidopsis* defense signaling network, required for resistance to the bacterial phytopathogen *Pseudomonas syringae* pv. *tomato* DC300^25,37,41,43,44^. As a mediator of the plant PM-CW continuum, NDR1 has been shown to play a role in regulating fluid movement across the membrane in response to pathogen infection, revealing that an increase in electrolyte leakage in the *ndr1* mutant following pathogen inoculation, as compared to WT Col-0^25,37^. As nutrient acquisition is the primary factor driving pathogen virulence, the observation of a reduction in the PM-CW continuum, resulting in a loss in cellular integrity and the ability to regulate the retention of nutrients, offers a plausible mechanism to explain the enhanced bacterial growth phenotype in the *ndr1* mutant. In total, these findings suggest that NDR1 is required for the maintenance of focal adhesion and the transduction of extracellular stimuli to intracellular signaling.

In the current study, we build upon our previous work to define the mechanism(s) through which NDR1 is required for abiotic and biotic stress signaling. Based on NDR1’s role in PM-CW adhesion, we hypothesized that NDR1 may also play a role in abiotic stress signaling, a mechanism which Singh et al.^14^ previously described for late embryogenesis abundant-14 (LEA14), a structural homolog of NDR1 and a known regulator of desiccation tolerance^25^. The hypothesis that NDR1 is likely associated with processes that converge on plant immunity and abiotic stress (i.e., drought) signaling is based on the described significance of water content during pathogen infections. Indeed, previous studies have shown that pathogens encounter inhibitory levels of osmotic potential once inside their host, and as part of their virulence response, deploy specific effector molecules which enhance colonization as a function of nutrient and water availability^45,46^. Here, we observed that under conditions of drought, RWC in the *ndr1* mutant decreased much faster than in WT Col-0 plants following water withholding. This decrease in RWC correlated with our previous observation of enhanced electrolyte leakage in the *ndr1* mutant^25^, supporting a role for NDR1 similar to that of the LEA family of proteins as functioning in processes that link biotic and abiotic stress activation through the perception of cell desiccation.

As a common denominator in the avoidance of drought stress and pathogen infection, the regulation of guard cell dynamics (i.e., stomata aperture) through ABA signaling represents a critical process in a plant’s response to environmental stress. At a regulatory level, previous data demonstrate that ABA metabolism and catabolism are modulated during host-pathogen interactions, and moreover, that increased levels of ABA drive host susceptibility to pathogen infection^47^. As a function of the role of NDR1 in this mechanism, data presented herein correlate a decline in RWC in the *ndr1* mutant with an increase in gene expression of the key regulators of ABA metabolism. As a regulatory point in describing the disconnect between increases in ABA biosynthesis (both mRNA and metabolite levels), our data also show that increases in ABA biosynthesis in the *ndr1* mutant did not prevent the *ndr1* mutant from rapidly declining into severe drought stress conditions, and ultimately, death.

We propose that the *ndr1* mutant is unable to perceive changes in ABA levels as an indicator of stress, and as a result, does not activate the appropriate signaling responses. In the case of abiotic stress, this translates into increased sensitivity to drought. In the case of biotic stress, this results in increased fluid loss during pathogen infection, leading to an enhanced disease susceptibility phenotype. NDR1 has long been associated with disease resistance signaling, including a key role in ETI^27,41,43,44^ and PTI^25^. We next investigated the link between drought, PTI, and NDR1 in stomatal-based signaling. To this end, we examined the role(s) of NDR1 in the regulation of stomata aperture, addressing both the abiotic (i.e., ABA) and biotic (i.e., PAMP) potential for NDR1-mediated signaling. We observed an abrogation in stomata function in the *ndr1* mutant through both an ABA-dependent manner, as well as through PTI-based immune signaling. These data are in agreement with a loss in sensitivity to ABA, as well as offer a possible mechanism supporting the enhanced disease susceptibility phenotype observed in the *ndr1* mutant^25,41,44^.

The activation of PM H^+^-ATPase(s) leads to hyperpolarization of the membrane and subsequent induction of inward ion channels, the result of which is stomatal opening (Fig. 8c). In contrast, inhibition of the PM H^+^-ATPase and subsequent stimulation of anion channel activation initiates membrane depolarization, resulting in the activation of outward rectifying K^+^ channels and stomatal closure^48,49^, less turgor pressure, and the abrogation of stomata closure. This data is further in agreement with recent work by Liu et al.^42^ which demonstrated that RIN4 modulates the activity of the PM H^+^-ATPase to regulate stomatal aperture during pathogen infection.

Based on the observation that a *RIN4*-overexpression line exhibits enhanced disease susceptibility and increased PM H^+^-ATPase activity, we asked what role NDR1 might play in this process. As a foundation for this question, we exploited previous work by Day et al.^43^ which identified an interaction between NDR1 and RIN4. While the significance of this interaction is not fully defined, we hypothesize that one consequence of NDR1’s association with RIN4 is to titrate the negative regulatory function of RIN4. In the case of the mechanism described here, we hypothesize that in the absence of NDR1 (i.e., *ndr1*), there is more unassociated RIN4 protein available to interact with the H^+^-ATPase. Such a mechanism is supported by the results described in Liu et al.^42^ using *RIN4* overexpressing plants. Indeed, the basic principle of RIN4-interacting proteins titrating the negative-regulatory potential has been previously proposed^42, 43, 50^, and while direct evidence describing the stoichiometry of RIN4-associated protein complexes is unknown, corollary data support this hypothesis.

In the current study, we demonstrate that NDR1 functions in signaling processes associated with both biotic and abiotic stress response pathways, a function we hypothesize represents NDR1’s role in the maintenance of cellular homeostasis during stress. Similar to the activity of mammalian integrin^51,52^, we hypothesize that NDR1 links the perception of perturbations to cellular homeostasis to the activation of specific signal transduction processes, including the response to drought and pathogen infection. As a next step in understanding the mechanism of NDR1’s function in stress perception, it will be important to determine how NDR1 activity functionalizes various stress perception responses, and in turn, modulates downstream signaling to quickly and specifically respond to these stresses. We posit that this activity is mediated in large part by the differential association of NDR1 with various key signaling components, such as RIN4 (immunity) and AHA5 (drought).

## Methods

### Plant materials and growth conditions

*Arabidopsis thaliana* and *Nicotiana benthamiana* plants were grown in a BigFoot series growth chamber (BioChambers) at 21°C under a 12 h light/12 h dark cycle with 60% relative humidity and a light intensity of 120 μmol photons m^−2^s^−1^. All *Arabidopsis* plants described in this study were in the Columbia-0 (Col-0) background. For seedling-based assays, *Arabidopsis* seeds were surface sterilized in 50% (v/v) bleach and 0.05% (v/v) Tween-20 for 20 min and washed 5 times with sterile distilled water. Seeds were germinated in 6-well plates containing ½-strength Murashige-Skoog (MS; Sigma-Aldrich) and 0.5% (w/v) MES (2-morpholinoethanesulfonic acid; Sigma-Aldrich), pH 5.7, for 4 d at 40 rpm in an orbital shaker at room temperature.

### Genome sequencing

Rosette leaves of a single 4-week-old *ndr1*/*35S∷NDR1* plant were collected and ground into a fine powder in liquid nitrogen. Genomic DNA was extracted using the Qiagen plant DNeasy kit and quantified with the Qubit 1x DNA BR assay kit. A DNA-seq library was constructed using 500 nanograms of genomic DNA using the Kapa HyperPlus prep kit according to the manufacturer’s protocol with a fragmentation time of 20 min and five PCR cycles. Libraries were size-selected at 300-600 bp using SPRI beads. The resultant library was sequenced on an Illumina MiSeq using paired end 75 bp mode to achieve ~10X genome coverage.

Raw reads were trimmed and quality-filtered using Trimmomatic v0.38^53^. To identify the transgene insertion site(s), we *de novo* assembled the genome of *ndr1*/*35S∷NDR1* plant using SOAPdenovo^54^. A k-mer size of 31 was used, and all other parameters were left at default settings. The *NDR1* overexpression insert was identified using BLASTn with the plasmid sequence as a query against the assembled contigs. Contigs with 95% or less of their contig length covered by insert sequences were extracted and manually inspected. Contigs containing both the plasmid and *Arabidopsis* genome sequences were used to identify the transgene insertion site by mapping the *Arabidopsis* genomic sequences to TAIR10 assembly.

### Seed germination assays

*Arabidopsis* seeds were surface sterilized as described above and plated on ½-strength MS media containing 0.5% (w/v) MES (pH 5.7) and 0.8% (w/v) agar containing ABA (Sigma-Aldrich), NaCl, mannitol (Sigma-Aldrich), and PEG6000 (Sigma-Aldrich). Fifty seeds were plated per plate. For seed stratification, plates were wrapped in aluminum foil and placed at 4°C for 4 d. After 4 d, plates were transferred to a BioChambers model GC8-2H growth chamber set at 20°C with a 12 h light/12 h dark cycle, ~60% relative humidity, and a light intensity of 120 μmol photons m^−2^s^−1^. For germination assays, radicle emergence was scored as germination. Experiments were repeated 5 times (independent biological replicates), with n = ~250 seeds for each genotype.

### Root length assay

Fifteen seedlings of WT Col-0, *ndr1*, and *ndr1/35S∷NDR1* plants were grown on ½-strength MS agar plates for 6 d at 22°C in a Percival chamber 36L5 (Geneva Scientific) under a 12 h light/dark cycle with a light intensity of 155 μmol photons m^−2^s^−1^. After 6 d, germinated seedlings were transferred onto ½-strength MS plates supplemented with increasing concentrations of ABA, mannitol, PEG 6000, and NaCl. Plates were oriented vertically to facilitate linear root growth. Root length was measured 6 d after transfer onto stress-inducing plates (i.e., 12 d post-germination). The average distance of root elongation was calculated from 4 independent biological replicates from 15 seedlings per genotype per experiment.

### Drought stress analysis

Prior to sowing seeds, pots containing water and soil were weight-adjusted across all planted genotypes and experiments to standard water content. A single plant was grown in each pot. For watering, the same volume of water was added to the flats containing potted plants every two-to-three days, as needed. Pot weights were measured every day to calibrate water used/lost for each experiment. For initiation of drought conditions at 3-weeks post-germination, flats containing plants, randomized across genotypes, were bottom flooded so that each pot was saturated with water (Day 0). Flats were randomly arranged in the growth chamber to minimize positional effects (e.g., airflow and light intensity). Plants were subjected to two watering regimes: 1) well-watered, wherein plants were watered three times weekly; and 2) drought stress (DS) conditions, wherein plants did not receive water. Leaf samples were harvested at 12, 14, 16, 17, 18, 19, and 21 d post-water withholding (dpw). Leaf samples for RNA extraction and hormone analysis were frozen in liquid nitrogen immediately and kept at −80°C until processing.

### Relative water content analysis

Relative water content (RWC) was evaluated using 4-to-5-week old plants (leaves) subjected to control and water-withholding conditions as follows: one fully expanded *Arabidopsis* leaf from each of 15 plants was harvested at 12, 14, 16, 17, 18, 19, and 21 dpw and immediately weighed to obtain the fresh weight (FW). Next, leaves were floated in sterile dH_2_O in a 15 ml sterile tube for 24 h at 4°C. After 24 h, the turgid weight (TW) of leaves was measured. Sampled leaves were then dried in an oven at 65°C for 24 h, removed to room temperature for 15 min, and the dry weight (DW) was measured. RWC was calculated using the formula:

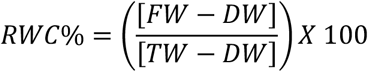

### Electrolyte leakage analysis

Electrolyte leakage analysis was performed on leaves from 4-to-5-week old plants subjected to control and drought stress conditions as follows: one fully expanded *Arabidopsis* leaf from each of 15 plants was harvested at 12, 14, 16, 17, 18, 19, and 21 dpw and immediately placed into a 15 ml sterile tube. Next, 4 ml of sterile dH_2_O was added to each tube and the samples were incubated overnight at 4°C. After overnight incubation, the samples were removed from 4°C and were acclimated to room temperature (25°C, ~1 h). Electrolyte readings (μS cm^−1^) were recorded using a SevenCompact S230 conductivity meter (Mettler Toledo), and after collection, samples were boiled for 30 min. After boiling, all samples were cooled to room temperature (25°C; ~1 h), and a final reading was collected. Percent electrolyte leakage was calculated using the following formula:

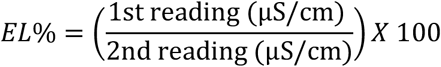

### Phytohormone analysis and quantification

Quantification of phytohormones was performed as previously described^55^, with minor modifications. In brief, 100 mg fresh weight (2-3 leaves, 4-to-5 week-old plants) were flash frozen in liquid nitrogen and stored at −80°C until processing. For extraction, frozen tissue was ground using a TissueLyser II (Qiagen) and incubated on a rocking platform at 4°C for 24 h in extraction buffer (80:20 v/v HPLC-grade methanol:water with 0.1% formic acid (v/v), 0.1 g l^−1^ butylated hydroxytoluene). Samples were centrifuged at 12,000 x *g* for 10 min at 4°C and resultant supernatants were collected and were filtered through a 0.2 mm PTFE (polytetrafluoroethylene) membrane (Millipore). Abscisic acid (ABA)-d_6_ (gift from Dr. Daniel Jones, MSU) served as an internal standard. Injections of plant extracts (10 μl per injection) were separated on a fused core Ascentis Express C18 column (2.1 × 100 mm, 2.7 μm; Supelco) installed in the column heater of an Acquity Ultra Performance Liquid Chromatography (UPLC) system (Waters Corporation). A gradient of 0.15% aqueous formic acid (solvent A) and methanol (solvent B) was applied in a 5-min program with a mobile phase flow rate of 0.4 μl min^−1^ as follows: 0 to 0.5 min hold at 99% A/1% B, ramp to 70% B at 3 min, ramp to 100% B at 3.01 min, hold at 100% B to 3.5 min, return to 99% A at 3.51 min and hold at 99% A to 5 min. The column was maintained at 40°C and interfaced to an Acquity TQ-D LC-MS/MS system equipped with electrospray ionization and operated in negative ion mode. The capillary voltage and extractor voltage were set at 2.5 kV and 3 V, respectively. The flow rates of cone gas and desolvation gas were 40 and 800 L h^−1^, respectively. The source temperature was 130°C and the desolvation temperature was 350°C. Collision energies and source cone potentials were optimized for each compound using QuanOptimize software (Waters Corporation). Peak areas were integrated, and the analytes were quantified based on standard curves generated from peak area ratios of analytes. Data acquisition and processing was performed using Masslynx 4.1 software (Waters Corporation). Analytes were quantified by converting peak area to phytohormone concentration (nM) per gram of dry weight of leaf tissue using a standard curve specific to each compound.

### Stomata aperture measurements

Leaf punches (0.25 cm^2^) from 4-to-5-week old plants were floated on opening buffer for 2-3 h at room temperature (25°C) before assaying. After acclimation, control treatments were imaged to establish a baseline for stomatal guard cell aperture. For evaluation of abiotic and biotic treatments, leaf punches were transferred to stress-inducing conditions (e.g., ABA, PEG6000, Mannitol, and H_2_O_2_) for 1 h. After 1 h, leaf punches were visualized using an Olympus IX71 microscope at 100X magnification and images were collected using an attached DP71 camera (Olympus). For each treatment, ~25 stomata were imaged for each plant genotype evaluated. Images were analyzed using ImageJ and the data were expressed as a width by length ratio (W/L). Experiments were repeated 3 independent times. Statistical significance was determined using a one-way ANOVA followed by Tukey’s post-test.

### Agrobacterium transient expression and co-immunoprecipitation assays

The CDS of *AHA1*, *AHA2*, *AHA5*, and *mCherry* (as the negative control) were cloned to binary vector pEarlygate201 with an HA tag, and NDR1 is cloned to pEarlygate202 containing a FLAG tag. We optimized *AHA5* codon using Invitrogen GeneArt GeneOptimizer to synthesize the CDS of *AHA5*.

Co-immunoprecipitation (co-IP) experiments were performed as previously described^25^, with slight modifications. *A. tumefaciens* strains expressing epitope-tagged constructs of *NDR1*, *AHA1, AHA2*, and *AHA5* fused to a 35S cauliflower mosaic virus promoter (constitutive expression) were used for co-IP assays. For transient expression in *Nicotiana benthamiana*, *A. tumefaciens* GV3101 strains harboring expression constructs of interest were pre-incubated at room temperature (ca. 22°C) in induction media (10 mM MES pH 5.6, 10 mM MgCl_2_, 150 mM acetosyringone [Sigma-Aldrich]) for 2 h before hand-infiltration into 5-week-old *N. benthamiana* leaves using a 1-ml needleless syringe at a final concentration of 4 × 10^8^ cells ml^−1^ for each construct. Inoculated plants were kept at 22°C for 40 h (12 h light/12 h dark). Infiltrated leaves were incubated at room temperature (ca. 25°C) under continuous light for 40 h. After incubation, 15-1 cm^2^ leaf discs were harvested, flash-frozen in liquid nitrogen, and stored at −80°C until processed. Frozen samples were ground into a fine powder in liquid nitrogen using a mortar and pestle. Once pulverized, the powder was transferred to a chilled mortar and pestle and further homogenized in extraction buffer (50 mM (4-(2-hydroxyethyl)-1-piperazineethanesulfonic acid) [HEPES] pH 7.5, 50 mM NaCl, 10 mM ethylenediaminetetraacetic acid [EDTA], 0.2% (v/v) Triton X-100, and 1 cOmplete, mini, EDTA-free protease inhibitor cocktail tablet [Sigma; 1 tablet/50 ml homogenization buffer]). Following homogenization, samples were transferred using a pipette to sterile 1.5 ml Eppendorf tubes. Samples were centrifuged at 15,000 x *g* for 20 min at 4°C, and the resultant supernatant (1 ml per tube) was transferred to 1.5 ml tubes. Five μl of the designated monoclonal antibody was added to each tube; an input control containing no antibody was also prepared. Following incubation at 4°C for 1 h (mixing end-over-end), 50 μl of pre-washed protein-G sepharose-4 fast flow (GE Life Sciences) was added to each tube and tumbled end-over-end for 4 h at 4°C. After 4 h, immunocomplexes were pelleted by centrifugation (5000 x *g*) and were washed 3 times in wash buffer (50 mM HEPES pH 7.5, 50 mM NaCl, 10 mM EDTA, 0.1% (v/v) Triton X-100, and 1 cOmplete mini EDTA-free protease inhibitor cocktail tablet [1 tablet/50 ml homogenization buffer]). After the final wash, immunocomplexes were pelleted by centrifugation (5000 x *g*), the wash buffer was removed by pipetting, and the pelleted samples resuspended in 50 μl of 3x Laemmli buffer [4% (w/v) SDS, 20% (v/v) glycerol, 0.004% (w/v) bromophenol blue, 0.125 M Tris-Cl pH 6.8, and 10% (w/v) DTT (dithiothreitol)]. Samples were boiled for 5 min, cooled on ice (1 min), and 10 μl of the denatured samples was separated by SDS-PAGE. Following SDS-PAGE, samples were transferred to nitrocellulose (GE Healthcare Life Sciences). Signals were detected using the Super Signal West Pico chemiluminescent substrate (ThermoFisher), with anti-HA-HRP (1:1000; Roche), anti-Flag-HRP (1:1000; Sigma), or anti-cMyc-HRP (1:1000; AbCam).

### Immunoprecipitation and tandem mass spectroscopy

To identify NDR1-interacting proteins, leaves from 5-week-old transgenic *Arabidopsis* plants expressing native promoter driven T7-NDR1 (i.e., *ndr1*/*P_NDR1_∷T7-NDR1*;^25^) were harvested as described above (co-immunoprecipitation assays). Total protein extracts were incubated with 5 μl T7 antibody (EMD Millipore; Cat # 69522) for 4 h, tumbling end-over-end, at 4°C. Following incubation, 50 μL of protein-G sepharose-4 fast flow (GE Life Sciences) (pre-washed and suspended in: pre-washed (50 mM HEPES pH 7.5, 50 mM NaCl, 10 mM EDTA, 0.1% (v/v) Triton X-100, and 1 cOmplete mini EDTA-free protease inhibitor cocktail tablet [1 tablet/50 ml homogenization buffer]) was added to each sample and tumbled end-over-end at 4°C for 16 h. Immunoprecipitation reactions were washed 3 times in wash buffer (50 mM HEPES pH 7.5, 50 mM NaCl, 10 mM EDTA, 0.1% (v/v) Triton X-100, and 1 cOmplete mini EDTA-free protease inhibitor cocktail tablet [1 tablet/50 ml homogenization buffer]), and suspended in 3x Laemmli buffer. The reaction was boiled for 5 min, and samples were separated by SDS-PAGE (15% Tris-Bis).

Resolved gels were stained with CBB (Coomassie brilliant blue), and the stained protein band corresponding to the expected molecular weight of T7-NDR1 was excised and subjected to in-gel digestion according to previously described methods^56^, with slight modification. Briefly, excised gel bands were dehydrated in 100% acetonitrile (ACN) and incubated with 10 mM DTT in 100 mM ammonium bicarbonate (pH 8.0) at 56°C for 45 min. After 45 min, samples were dehydrated again and incubated in the dark with 50 mM iodoacetamide in 100 mM ammonium bicarbonate for 20 min. Gel bands were then washed with ammonium bicarbonate and dehydrated again. Sequencing grade modified trypsin was prepared at a concentration of 0.01 μg μl^−1^ in 50 mM ammonium bicarbonate, and 50 μl of trypsin was added to each sample such that the gel was completely submerged. Samples were incubated overnight at 37°C. After incubation, peptides were extracted from the gel by water bath sonication in a solution of 60% (v/v) ACN/1% (v/v) trichloroacetic acid (TCA) and vacuum dried to ~2 μl. Peptide samples were then re-suspended in 2% (v/v) ACN/0.1% (v/v) trifluoroacetate (TFA) to 20 μl. From this, 5 μl was injected onto a Thermo Acclaim PepMap RSLC 0.075 mm x 20 mm C18 trapping column and washed for 5 min using a Thermo EASYnLC. Peptides were eluted onto a Thermo Acclaim PepMap RSLC 0.075 mm x 500 mm C18 analytical column over 35 min with a gradient of 2% B to 40% B over 24 min, ramping to 100% B at 25 min and held at 100% B for the duration of the run (Buffer A = 99.9% water/0.1% (v/v) formic acid, Buffer B = 80% ACN/0.1% (v/v) formic acid/19.9% (v/v) water).

Eluted peptides were sprayed into a Q-Exactive HF-X mass spectrometer (ThermoFisher) using a FlexSpray spray ion source. Survey scans were taken in the Orbitrap (45,000X resolution, determined at *m/z* 200) and the top twenty ions in each survey scan were then subjected to automatic higher energy collision induced dissociation (HCD) with fragment spectra acquired at 7,500X resolution. Resultant tandem mass spectrometry (MS/MS) spectra were converted to peak lists using Mascot Distiller (ver. 2.7.0; www.matrixscience.com) and the output was searched against all entries in the TAIR protein sequence database (www.arabidopsis.org; ver. 10). All entries in the UniProt *E. coli* protein sequence database (downloaded from www.uniprot.org, 2017-11-01), appended with common laboratory contaminants (downloaded from www.thegpm.org, cRAP project), were scanned using Mascot (ver. 2.6). The Mascot output was analyzed using Scaffold Q+S (ver. 4.8.4; www.proteomesoftware.com) to probabilistically validate protein identifications. Assignments validated using the Scaffold 1% FDR (false discovery rate) confidence filter were considered true.

### Biomolecular fluorescence complementation assays

The CDS of *AHA1*, *AHA2*, *AHA5* (codon optimized, described above), and *WRKY40* (as the negative control) were cloned into binary vector pBH1096 (2X35S∷Gateway∷HA-cEYFP) for cEYFP fusion, and *NDR1* was cloned into pBM1089 (2X35S∷myc-nEYFP∷Gateway) for nEYFP fusion. A mixture of Agrobacterium strains expressing nEYFP- and cEYFP-fusion constructs was hand-infiltrated into *N. benthamiana* using the same approach as described above for co-IP assays. Leaf samples were harvested at 2 dpi and were imaged using a confocal microscope system (Olympus FluoView 1000). To accurately compare samples with positive/negative fluorescence, all images were captured using the same laser intensity, dwell time, and PMT voltage. The same linear transformation, abiding by academic criterion of biological imaging, was applied to the LUTs of all images to enhance contrast.

### Plasma membrane H^+^-ATPase activity assay

Plasma membrane isolation was performed according to previously published methods^57^. In brief, 30 g of 5-week-old *Arabidopsis* leaves were homogenized in 100 ml ice-cold homogenization buffer containing 0.33 M sucrose, 50 mM Hepes-KOH (pH7.5), 10% glycerol (wt/vol), 0.2% BSA (wt/vol), 5 mM DTT (dithiothreitol), 5 mM EDTA, 5 mM ascorbate, 0.6% polyvinylpyrrolidone, 1 mM PMSF (phenylmethylsulfonyl fluoride) and cOmplete EDTA-free protease inhibitor cocktail tablet [1 tablet/100 ml homogenization buffer]). The homogenate was passed through five layers of Miracloth to remove debris and was then centrifuged at 18,000 x *g* for 20 min at 4°C. A microsomal pellet was isolated by ultracentrifugation at 120,000 x *g* for 30 min at 4°C and resuspended in a buffer containing 0.33 M sucrose, 3 mM KCl, 0.1 mM EDTA, 1 mM DTT, 1 mM PMSF, 5 mM potassium phosphate (pH7.8) and cOmplete EDTA-free protease inhibitor cocktail tablet [1 tablet/100 mL buffer]). A plasma membrane-enriched fraction was purified from the microsomal fraction by two-phase partitioning^58^. The resuspended microsomal fraction was added to a centrifuge tube containing dextran T500 and polyethylene glycol (PEG) 3350 in 0.33 M sucrose, 3 mM KCl and 5 mM potassium phosphate (pH 7.8). The contents were mixed by inverting 20-25 times and centrifuged in a swinging bucket rotor at 1000 x *g* for 10 min at 4°C. The PEG upper phase was removed, diluted 5-fold in suspension buffer (0.33M sucrose, 20 mM Hepes-KOH (pH 7.5), 10% glycerol (wt/vol), 0.1% BSA (wt/vol), 2 mM DTT (dithiothreitol), 0.1 mM EDTA, cOmplete EDTA-free protease inhibitor cocktail tablet [1 tablet/100 ml homogenization buffer]) and centrifuged at 120,000 x *g* for 45 min. The resultant plasma membrane pellet was solubilized in suspending buffer (0.33 M sucrose, 20 mM Hepes-KOH (pH 7.5), 10% glycerol (wt/vol), 0.1% BSA (wt/vol), 2 mM DTT (dithiothreitol), 1 mM EDTA, cOmplete EDTA-free protease inhibitor cocktail tablet [1 tablet/100 ml homogenization buffer]) to a final concentration between 2-4 mg ml^−1^. H^+^-pumping activity was detected by a decrease in ACMA absorbance at 495 nm^59^.The assay buffer contained 20 mM MES-KOH (pH 7.0), 4 mM ATP-Na_2_, 140 mM KCl, 100 μM ACMA (9-amino-6-chloro-2-methoxy acridine), 0.05% Brij 58, and 50 μg of plasma membrane protein in a total volume of 1 ml. Plasma membrane-enriched samples were pre-incubated at 25°C for 5 min in assay buffer. The assay was initiated by the addition of 4 mM MgSO_4_ and the absorbance (*A*_495nm_) was measured over a time interval from 4 to 5 min. Each experiment was repeated 2-3 times with independent plasma membrane preparations.

### Quantitative real-time PCR (qPCR)

Total RNA was extracted from the leaves of 4-to-6-week old plants using the RNeasy Plant Mini Kit (Qiagen). RNA concentrations were quantified using a Qubit RNA BR Assay Kit (ThermoFisher) in combination with a Qubit 4 Fluorometer (ThermoFisher). First-strand cDNA was synthesized from 1 μg total RNA using the SuperScript™ III first-strand synthesis system (ThermoFisher). Quantitative real-time PCR (qRT-PCR) was performed using the Applied Biosystems 7500 Real-Time PCR instrument (ThermoFisher) using the SYBR Select qPCR master mix (ThermoFisher). Ubiquitin (*UBQ10*) served as an internal control for amplification. The abundance of target gene transcripts was normalized to *UBQ10* according to the 2^−ΔΔCT^ method^60^. All qPCR DNA primers utilized in this study are listed in Supplementary Table 1.

### TaqMan gene expression assay

Leaves from 3-week-old wild type (WT) Columbia-0 (Col-0), mutant, and complementation lines (i.e., *ndr1, ndr1/35S∷NDR1, aha5-2, and aha1-9*), were collected and immediately frozen in liquid nitrogen. Total RNA was extracted using the RNeasy Plant Mini kit (Qiagen). cDNA was synthesized with Maxima H Minus First Strand cDNA Synthesis Kit (Thermo-Fisher). Confirmation of the *aha5-2* mutant line was performed using TaqMan Gene expression Assay kits (Applied Biosystems) for HA5 (Assay ID At02274124_g1) and *UBQ10* (Assay ID At 02353385_S1) according to manufacturer’s protocol in ABI StepOnePlus thermocycler v2.3 (Applied Biosystems). The relative levels of each mRNA transcript were calculated and normalized to the expression of the *UBQ10* gene.

### Library preparation and sequencing

mRNA RNAseq libraries were prepared using the Illumina TruSeq mRNA library preparation kit from three biological repeats of each timepoint of well-watered or water-withholding conditions of WT Col-0, *ndr1*, *ndr1*/*P_NDR1_∷T7-NDR1*^25^, and *ndr1*/*35S∷NDR1*^27^ plants by the Michigan State Research Technology and Support Facility (RTSF). Multiplexed samples were pooled and sequenced on the Illumina HiSeq 4000 (single end 50 bp mode). Base calling was performed using the Illumina Real Time Analysis (RTA; v2.7.7), and the output of RTA was demultiplexed and converted to FastQ format using Illumina Bcl2fastq (v2.19.1).

### Analysis of RNA-sequencing

The adapter sequences and low-quality bases (q < 10) were trimmed by Trimmomatic^53^. Resultant cleaned reads were mapped to the TAIR10 reference genome using HISAT2^61^ Mapped read counts for each gene were generated using the HTseq^62^ command, and differentially expressed genes (DEG) were analyzed using DESeq2^63^. For the identification of DEGs, a threshold of *P*-value < 0.01 and at least a log_2_ fold change ≥ |1| was used as a criterion in this study. Gene ontology (GO) term enrichment analysis using Fisher test with a significance Level < 0.05 was performed by using AgriGO v2^64^.

Coexpression network analysis was performed using the R package WGCNA^65^. Normalized read counts of WT Col-0, the *ndr1* mutant, and the *NDR1*-overexpression line at 0, 12, 14 16, 17 18, 19, and 21 day timepoints under water-withholding treatment from DESeq2 were used for constructing a singed hybrid network. A matrix of Pearson correlation between all pairs of 31319 genes was calculated. The adjacency matrix was then constructed by using a power of 18, a mergeCutHeight of 0.25, and module size greater than 30. Average linkage hierarchical clustering was applied to the topological overlap for grouping genes with highly similar coexpression relationships. The expression profiles of each module was summarized by module eigengene resulting in 15 modules. Gene ontology (GO) term enrichment analysis was performed using Fisher test with a significance level < 0.05, and minimum number of mapping entries of 5 was performed by using AgriGO v2.

## Supporting information

Supplemental Data Sets

## Data availability

Sequence data from this article can be found in the *Arabidopsis* genome initiative or GenBank/EMBL databases under the following accession numbers: *At3g20610 (NDR1), At2g18960 (AHA1), At4g30190 (AHA2), At2g24520 (AHA5), At5g67030 (ABA1), At1g52340 (ABA2), At3g14440 (NCED3), At5g52300 (RD29B), At2g27150 (AAO3)*, and *At1g30360 (ERD4)*.

The Illumina RNA-seq reads were deposited in BioProject under the BioProject ID: PRJNA595619.

## Additional information

**Supplementary Fig. 1 | Extended timecourse phenotypes of drought-induced plants shown in Figure 1.** Each photo is representative of at least 15 individual plants per timepoint.

**Supplementary Fig. 2 | Drought phenotype recovery.** Photos were taken at 2 d post-watering following 25 d of water withholding.

**Supplementary Fig. 3 | Genome sequence analysis of *ndr1*/*35S∷NDR1*.** Illustration of the *35S∷NDR1* insertion site on chromosome 1.

**Supplementary Fig. 4 | ABA-induced inhibition of seed germination.** Fifty seeds were plated on a single Petri dish, per experiment. Radicle emergence was scored as germination. Varying concentrations of ABA were dissolved in MS media. **a,** 1 μM and **b,** 2 μM. Five independent biological replicates were carried out with n = ~250 seeds for each genotype.

**Supplementary Fig. 5 | Root elongation inhibition assays in the *ndr1* mutant in response to abiotic chemical stressors. a,** ABA, **b,** PEG6000, **c,** NaCl, and **d,** Mannitol. **e,** Root length of plants grown under normal conditions for 4 weeks in soil. Root length was measured at 6 days after transferring seedlings onto plates containing salt described above. The average of root elongation was measured from 4 independent biological replicates from 15 seedlings per genotype per experiment.

**Supplementary Fig. 6 | Relative mRNA level of *NDR1*.** *NDR1* mRNA accumulation in WT Col-0, the *ndr1* mutant, the *ndr1*/*P_NDR1_∷T7-NDR1* complementation line, and the *ndr1*/*35S∷NDR1* overexpression line under well-watered and drought conditions. Three biological repeats were performed (n=9), p<0.05.

**Supplementary Fig. 7 | mRNA expression levels of ABA related genes.** mRNA were measured at **a,** 16 day and **b,** 21 day post drought stress. Three biological repeats were performed (n=9), p<0.05.

**Supplementary Fig. 8 | Stomatal aperture during the diurnal cycle.** Plants were assayed every 2 h over a 24 h time period to analyze. Three biological repeats were performed (n=9), p<0.05.

**Supplementary Fig. 9 | Hormone accumulation during the 24 hours diurnal cycle**. **a,** ABA accumulation and **b,** SA accumulation. Three biological repeats were performed (n=9), p<0.05.

**Supplementary Fig. 10 | *Pst* DC3000 and *Pst* DC3000-AvrRpt2 *in planta* growth curves during drought stress.** Plants were subjected to drought stress and infected with **a,** the virulent pathogen *Pst*. DC3000 as well as the avirulent pathogen **b,** *Pst*-AvrRpt2. Plant were infiltrated with *Pst* DC3000 at 12 dpw. Three biological repeats were performed (n=9), p<0.05.

**Supplementary Fig. 11 | Principle component analysis (PCA) of gene expression values.** Gene expression values of both water withholding condition and well-watered condition during time course.

**Supplementary Fig. 12 | DEG information and GO terms under well-watered condition. a,** Expression profile of DEGs under well-watered conditions over the 12-21 day timecourse. **b,** Expression patterns of genes under well-watered conditions. **c,** GO terms of differential DEGs between *ndr1* vs WT Col-0 and *ndr1/35S∷NDR1* vs WT Col-0..

**Supplementary Fig. 13 | Expression patterns of transcription factors that response to drought stress.** Transcription factor mRNAs identified as up- or downregulated across all timepoints of the treatment.

**Supplementary Fig. 14 | Common up- or downregulated genes under well water and water withholding conditions. a,** The number of genes in *ndr1*/*35S∷NDR1* lines. **b,** The number of genes in ndr1 mutant. **c,** GO terms that are associated with genes identified from **a** and **b**.

**Supplementary Fig. 15 | Modules from coexpression profiles in WT Col-0, *ndr1*, and *ndr1/35S∷NDR1*.** Samples were collected at 0, 12, 14,16, 17, 18, 19 and 21 days under well-watered or water withholding conditions.

**Supplementary Fig. 16 | BiFC confirmed NDR1 associates with AHA1, AHA2, and AHA5.** Co-expression of n-YFP-NDR1 with H^+^-ATPase-cYFP proteins in *N. benthamiana* leaves. WRKY40-c-YFP was employed as a negative control.

**Supplementary Fig. 17** | **mRNA expression levels in 4 *Arabidopsis* genotypes. a,** *AHA1*, **b,** *AHA2*, **c,** *AHA5*, and **d,** *NDR1*.

## Acknowledgements

Research in the laboratory of B.D. is supported by a grant from the National Science Foundation Plant Biotic Interactions Program (IOS-1146128), the National Institute of General Medical Sciences (1R01GM125743), and the MSU Plant Resilience Institute (GR100125-Heat3). Research in the laboratory of R.V. was supported by startup funds from MSU. Research in the laboratory of I.B. is supported by the Scientific and Technological Research Council of Turkey (TUBITAK) in the context of 2219-Post Doctoral Fellowship Program. We are grateful to Mike Thomashow for critical comments and suggestions throughout the development of the study. We would like to thank Dan Jones (MSU) for assistance with phytohormone analysis and Doug Whitten (MSU) for proteomics analysis.

## Author contributions

Y.-J.L, H.C., A.C., I.B., P.S., and B.D. designed the experiments. Y.-J.L, H.C., A.C., P.L. and I.B., preformed the majority of the experiments and the analysis of the data, with H.S., S.S., C.M.W., H.S., P.S., B.V.B. and B.D. assisting. Y.-J.L., A.C., and B.D. wrote the article. All authors provided editorial comments during the development of this manuscript, and all authors approved the final manuscript before submission.

