## Supplemental Figures for "NDR1 and the Arabidopsis Plasma Membrane ATPase AHA5 are Required for Processes that Converge on Drought Tolerance and Immunity"

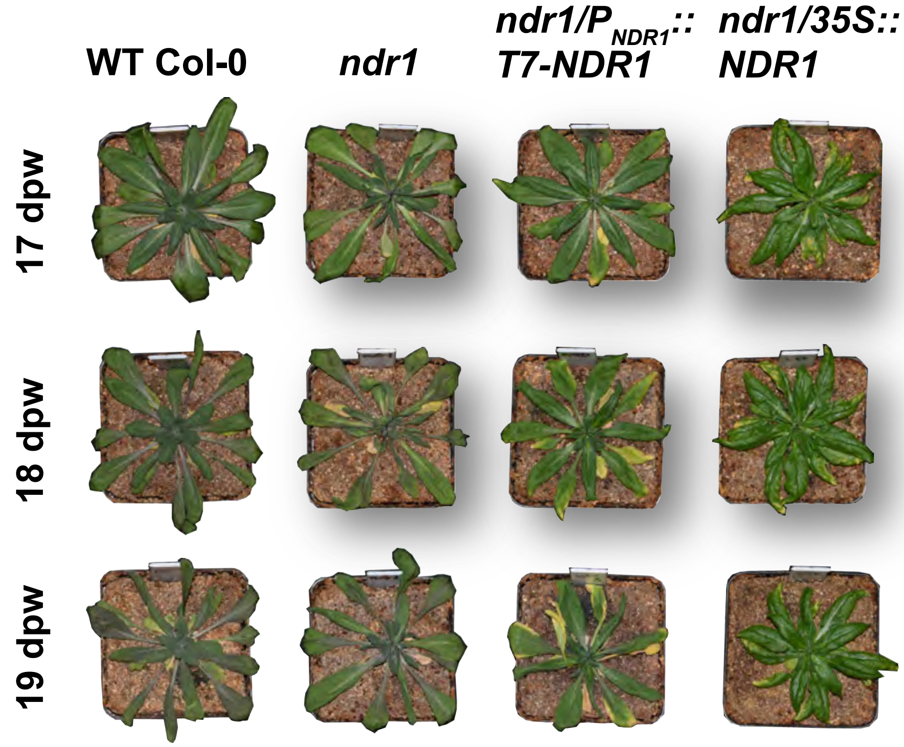


**Supplementary Fig. 1 | Extended timecourse phenotypes of drought-induced plants shown in Figure 1.** Each photo is representative of at least 15 individual plants per timepoint.

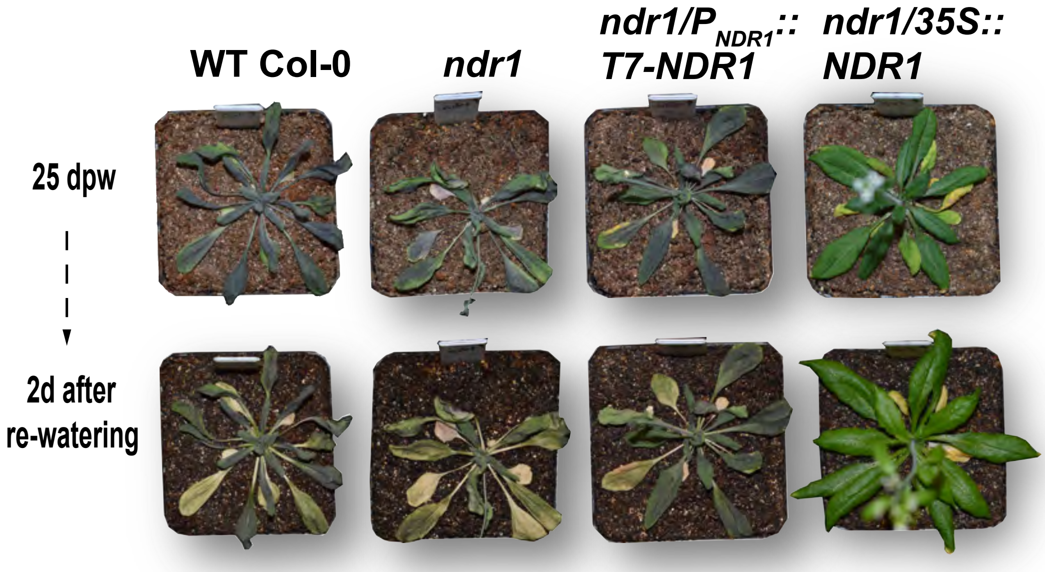


**Supplementary Fig. 2 | Drought phenotype recovery.** Photos were taken at 2 d post-watering following 25 d of water withholding.

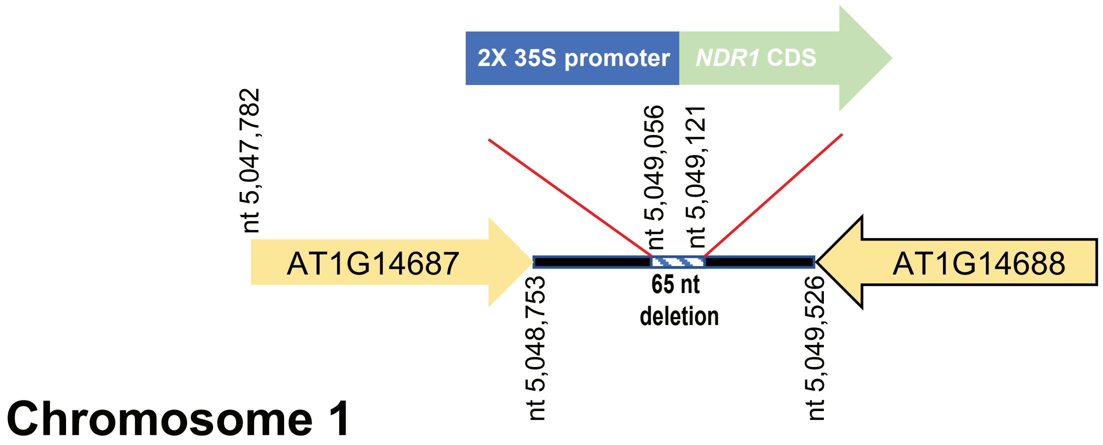


**Supplementary Fig. 3 | Genome sequence analysis of ndr1/35S::NDR1.**
Illustration of the 35S::NDR1 insertion site on chromosome 1.

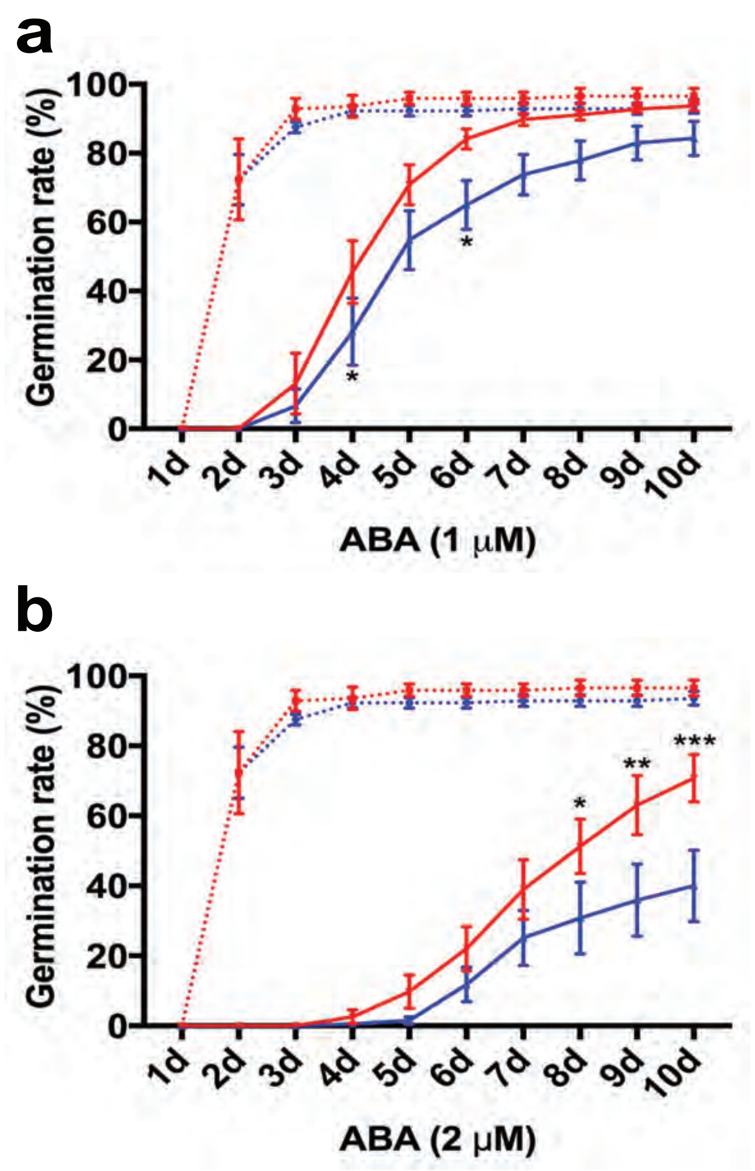


**Supplementary Fig. 4 | ABA-induced inhibition of seed germination.** Fifty seeds were plated on a single Petri dish, per experiment. Radicle emergence was scored as germination. Varying concentrations of ABA were dissolved in MS media. a, 1 uM and b, 2 uM. Five independent biological replicates were carried out with n = ~250 seeds for each genotype.

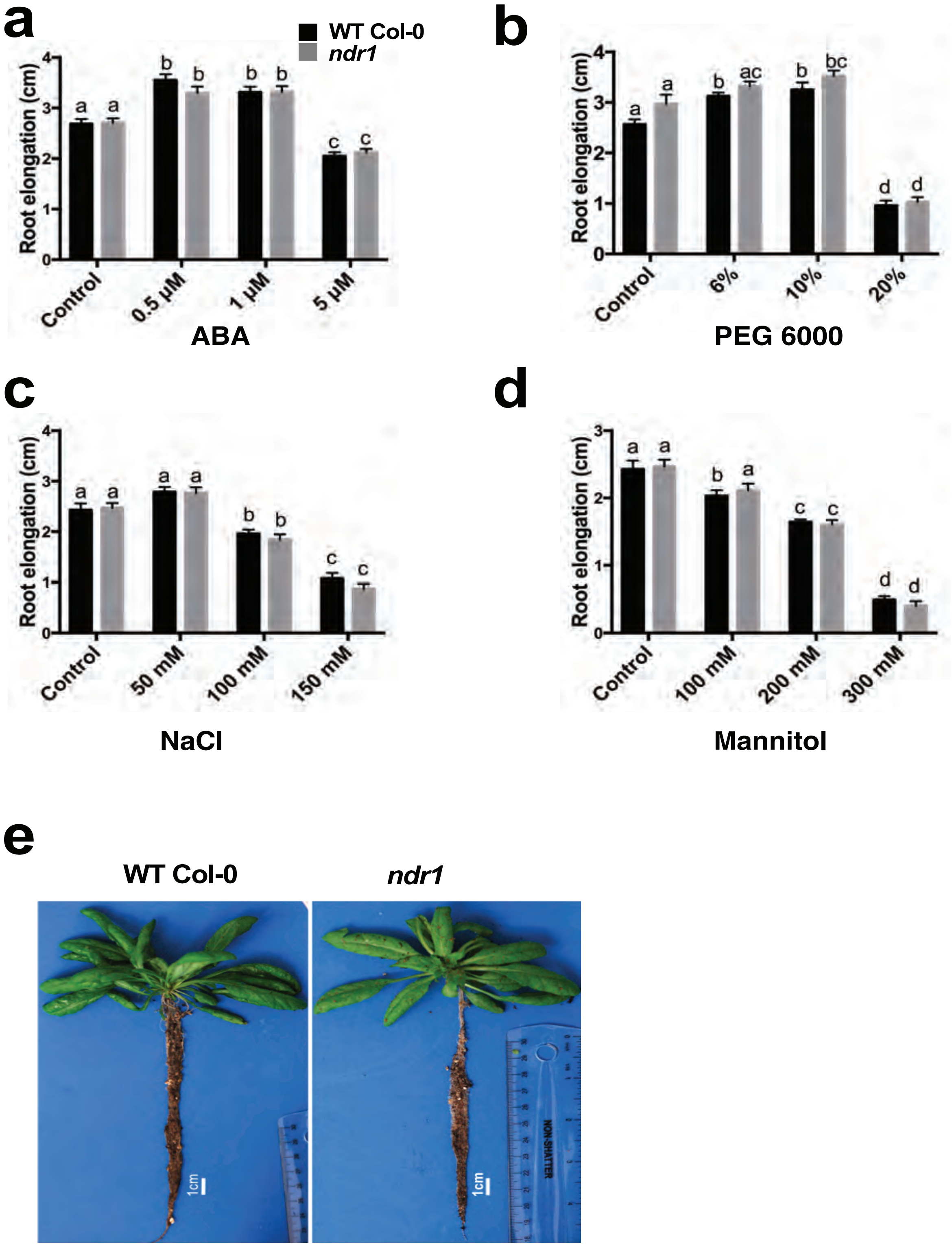


**Supplementary Fig. 5 | Root elongation inhibition assays in the ndr1 mutant in response to abiotic chemical stressors. a,** ABA, **b,** PEG6000, **c,** NaCl, and **d,** Mannitol. **e,** Root length of plants grown under normal conditions for 4 weeks in soil. Root length was measured at 6 days after transferring seedlings onto plates containing salt described above. The average of root elongation was measured from 4 independent biological replicates from 15 seedlings per genotype per experiment.

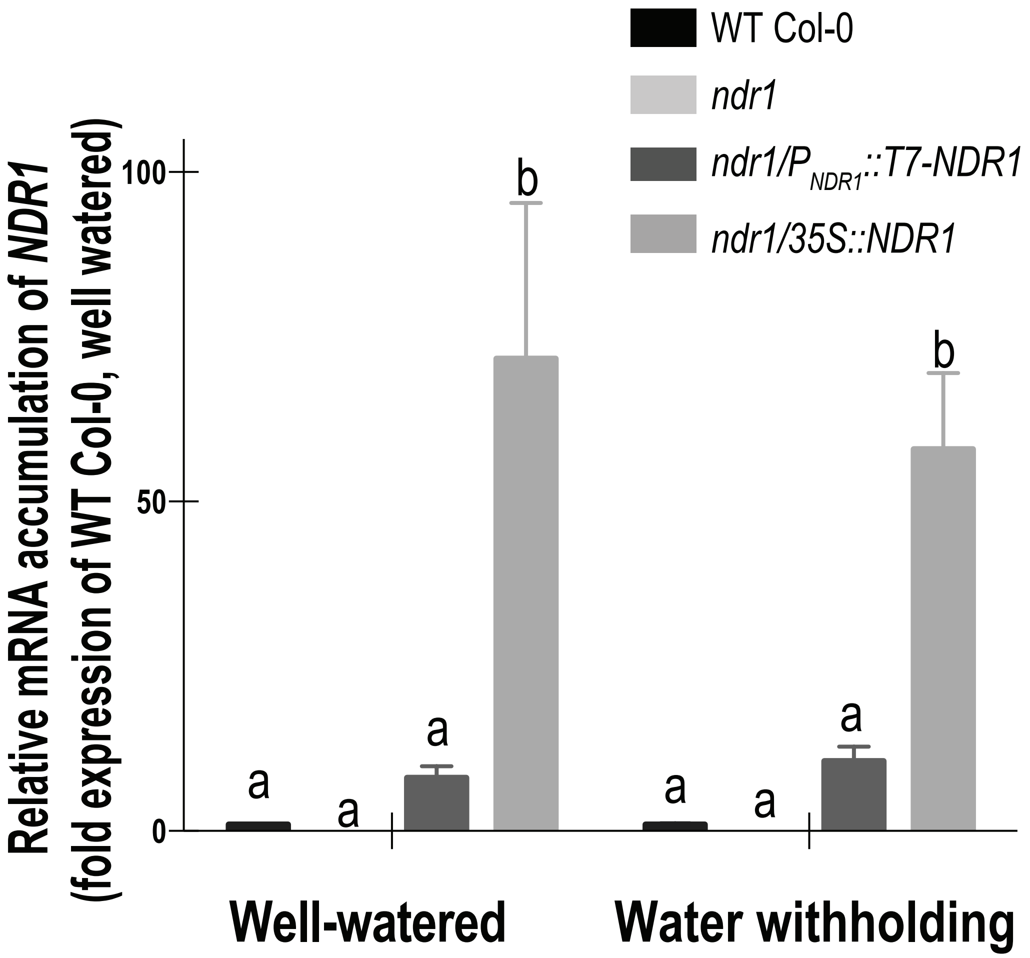


**Supplementary Fig. 6 | Relative mRNA level of NDR1.** NDR1 mRNA accumulation in WT Col-0, the ndr1 mutant, the ndr1/PNDR1::T7-NDR1 complementation line, and the ndr1/35S::NDR1 overexpression line under well-watered and drought conditions. Three biological repeats were performed (n=9), $p<0.05$.

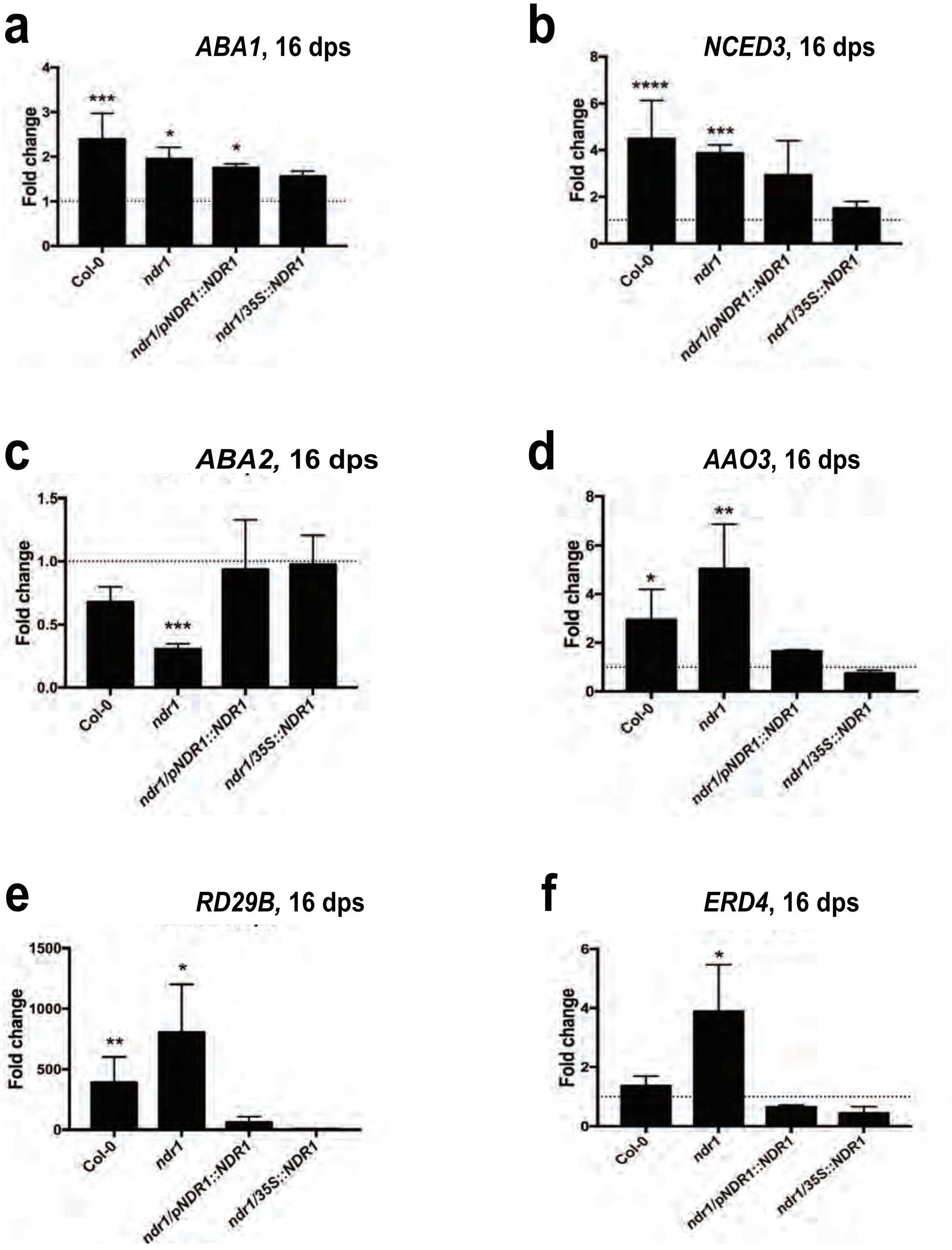


**Supplementary Fig. 7 | mRNA expression levels of ABA related genes.** mRNA were measured at a, 16 day and b, 21 day post drought stress. Three biological repeats were performed (n=9), p<0.05.

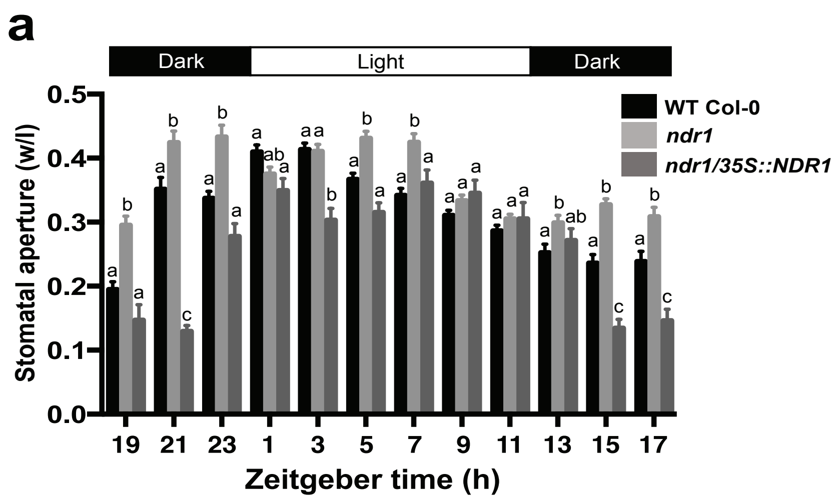


**Supplementary Fig. 8 | Stomatal aperture during the diurnal cycle.** Plants were assayed every 2 h over a 24 h time period to analyze. Three biological repeats were performed (n=9), p<0.05.

a

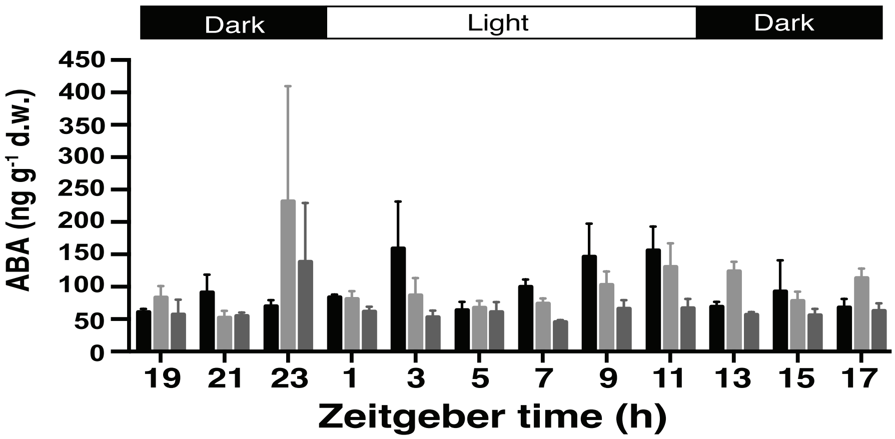


b

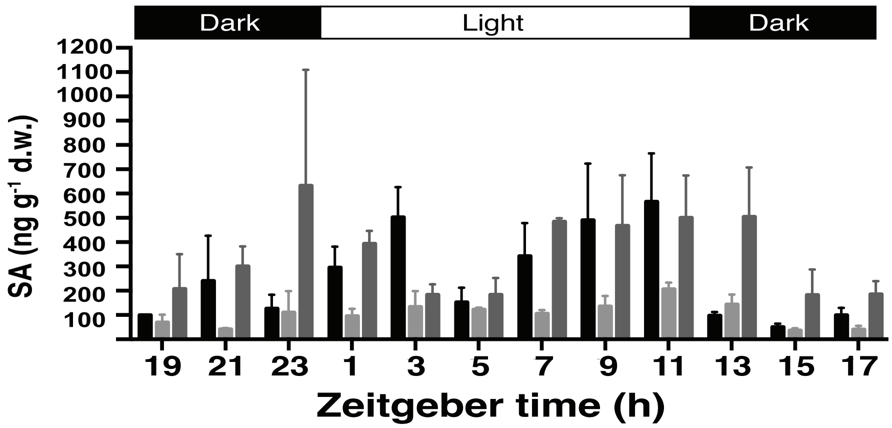


**Supplementary Fig. 9 | Hormone accumulation during the 24 hours diurnal cycle. a,** ABA accumulation and **b,** SA accumulation. Three biological repeats were performed (n=9), $p<0.05$.

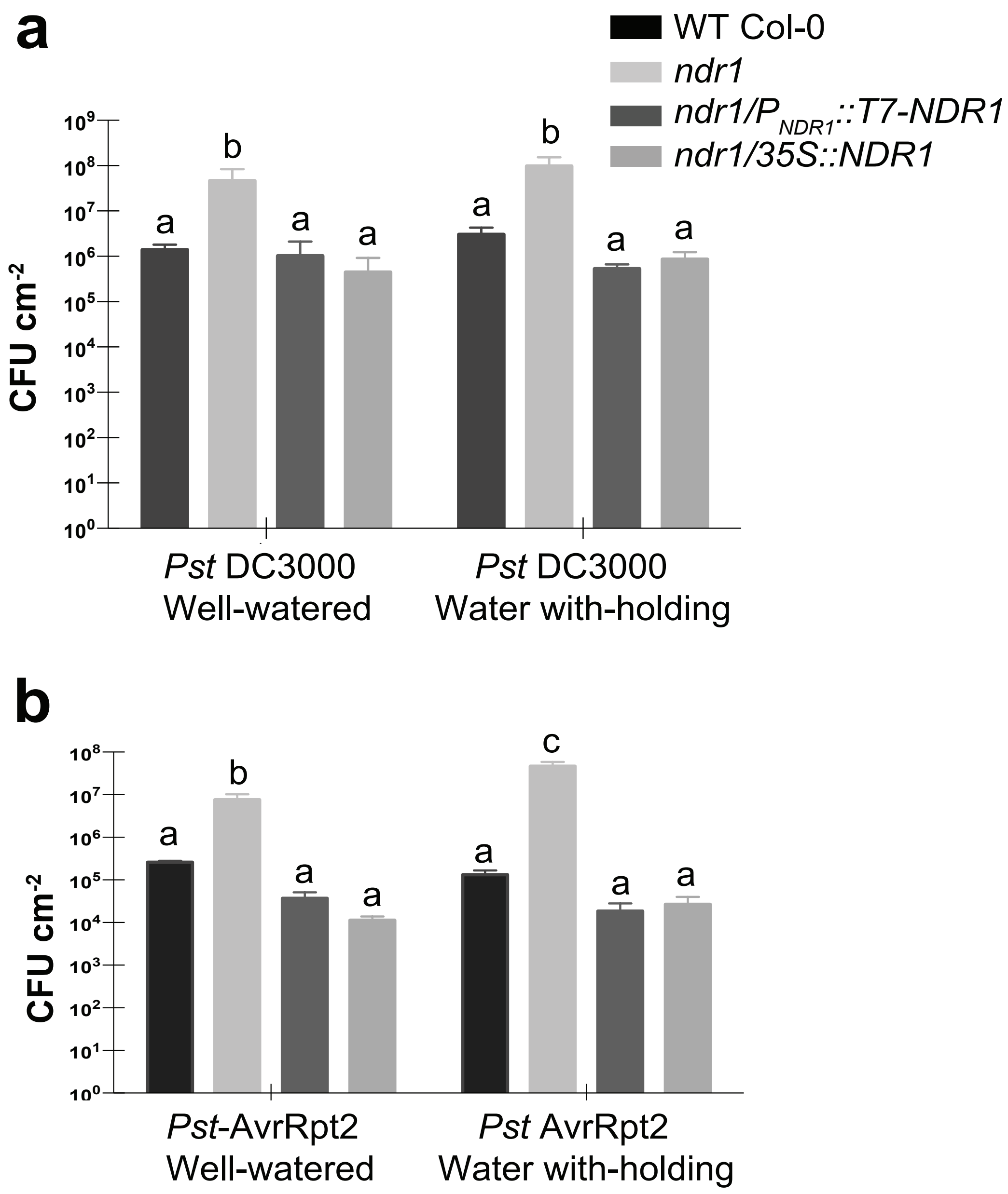


**Supplementary Fig. 10 | Pst DC3000 and Pst DC3000-AvrRpt2 in planta growth curves during drought stress.** Plants were subjected to drought stress and infected with **a,** the virulent pathogen Pst. DC3000 as well as the avirulent pathogen **b,** Pst-Avr-Rpt2. Plant were infiltrated with Pst DC3000 at 12 dpw. Three biological repeats were performed (n=9), $p<0.05$.

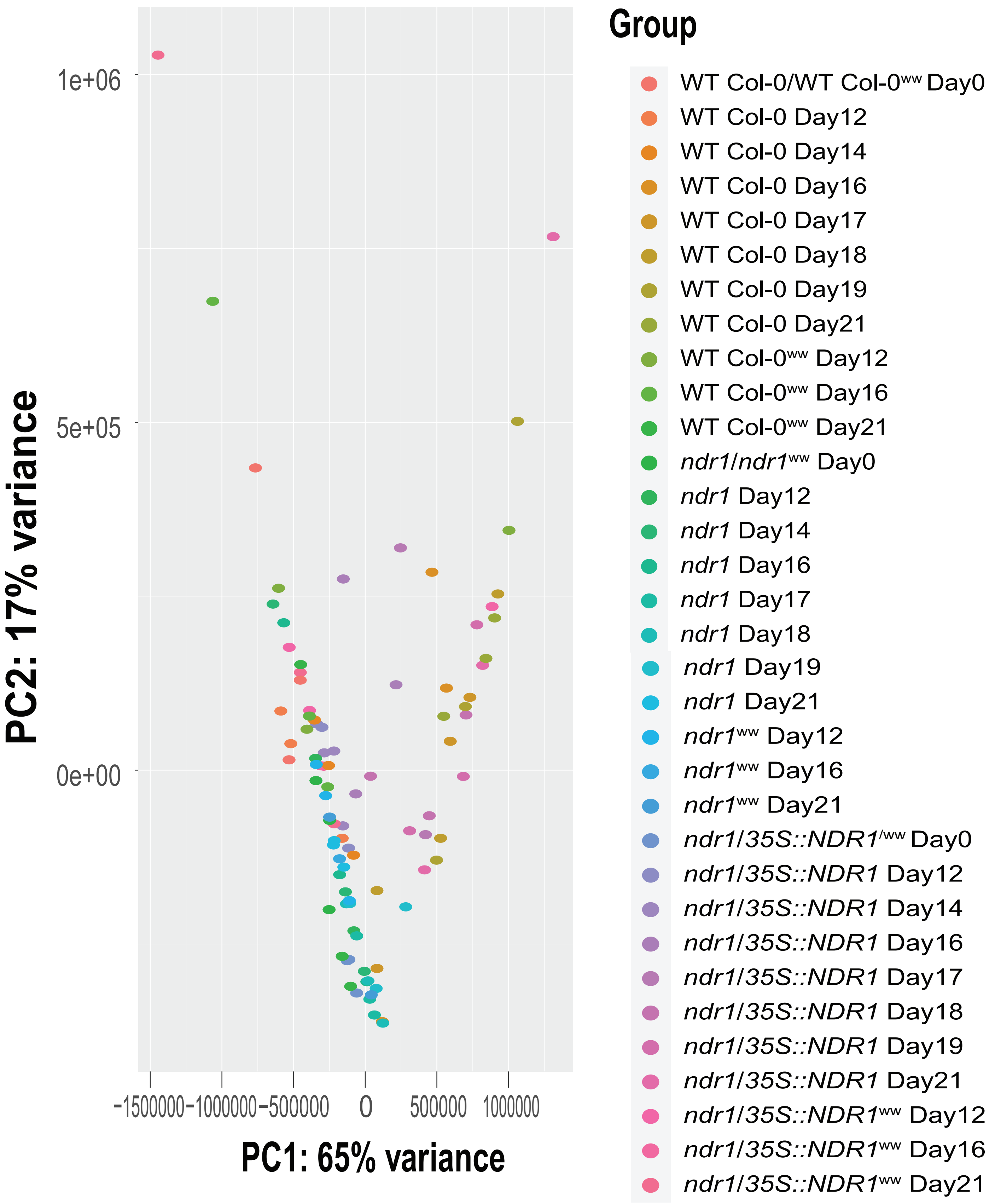


**Supplementary Fig. 11 | Principle component analysis (PCA) of gene expression values.** Gene expression values of both water with-holding condition and well-watered condition during time course.

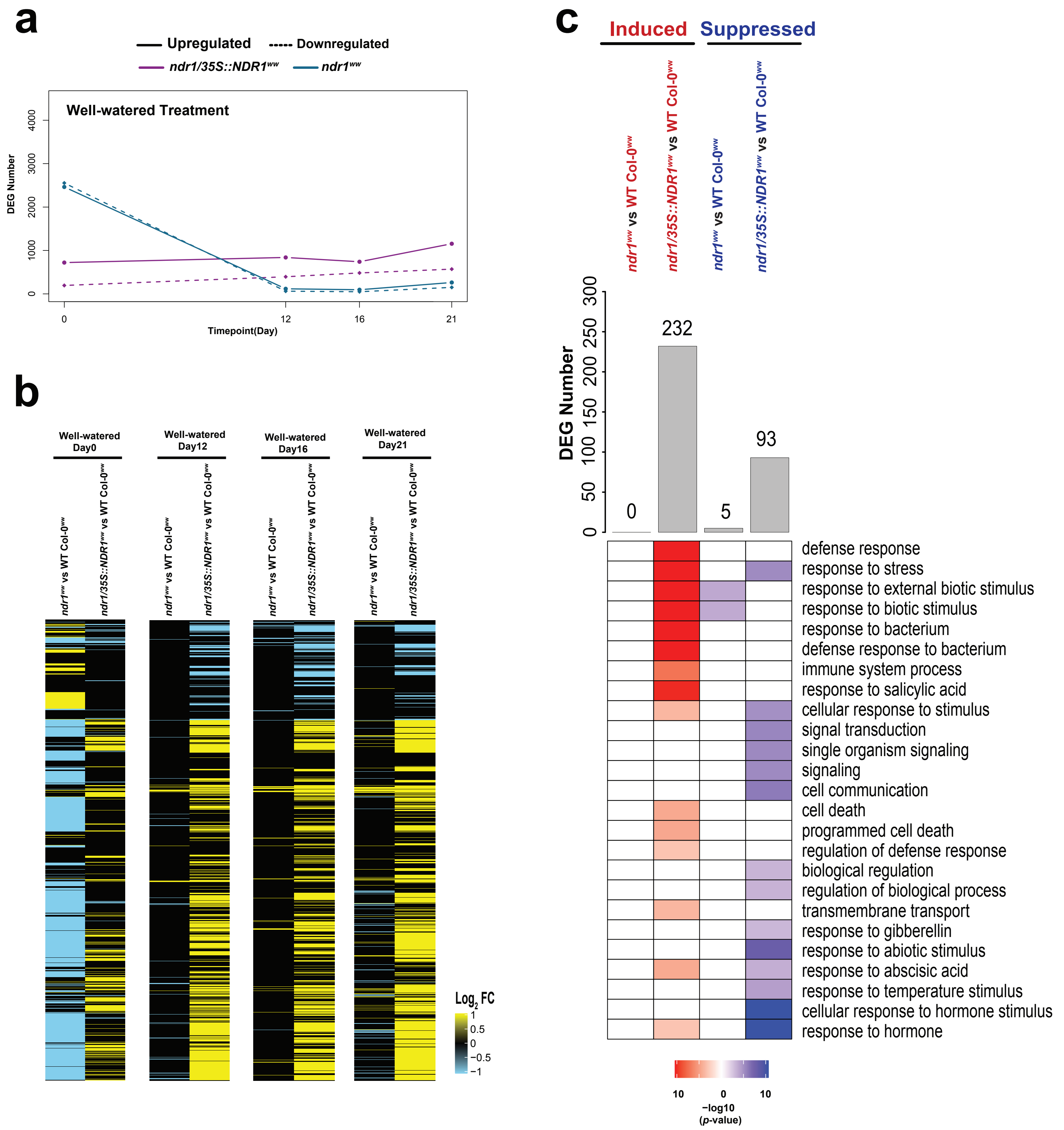


**Supplementary Fig. 12 | DEG information and GO terms under well-waterd condition. a,** Expression profile of DEGs under well-watered condition over the 12-21 day timecourse. **b,** Expression patterns of genes under well-wtered condition. **c,** GO terms of differential DEGs between *ndr1* vs WT Col-0 and *ndr1/35S::NDR1* vs WT Col-0.

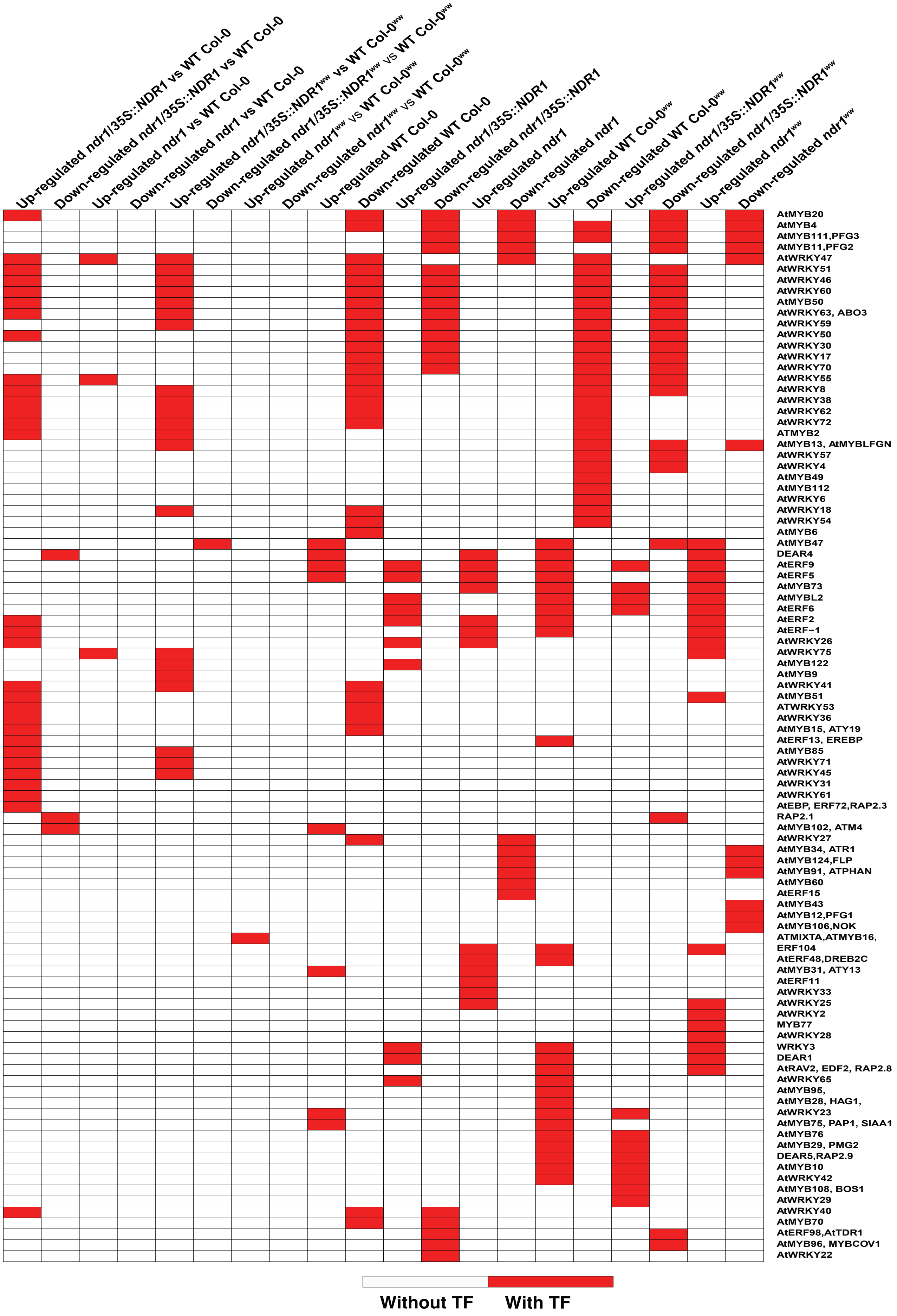


**Supplementary Fig. 13 | Expression patterns of transcription factors that response to drought stress.** Transcription factor mRNAs identified as up- or downregulated across all timepoints of the treatment.

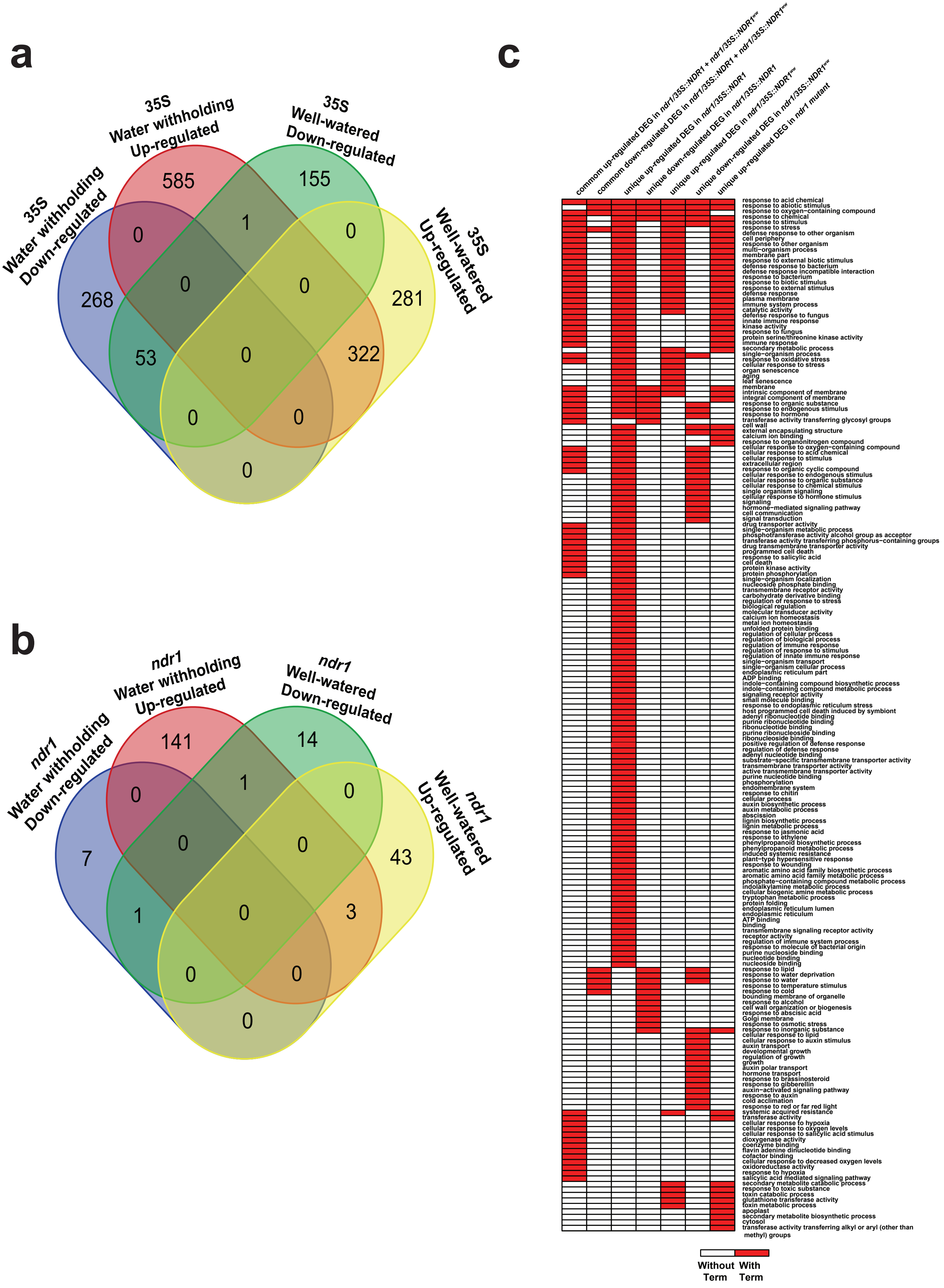


**Supplementary Fig. 14 | Common up- or downregulated genes under well water and water withholding conditions. a,** The number of genes in ndr1/35S::NDR1 lines. **b,** the number of genes in ndr1 mutant. **c,** GO terms that are associated with genes identified from **a** and **b**.

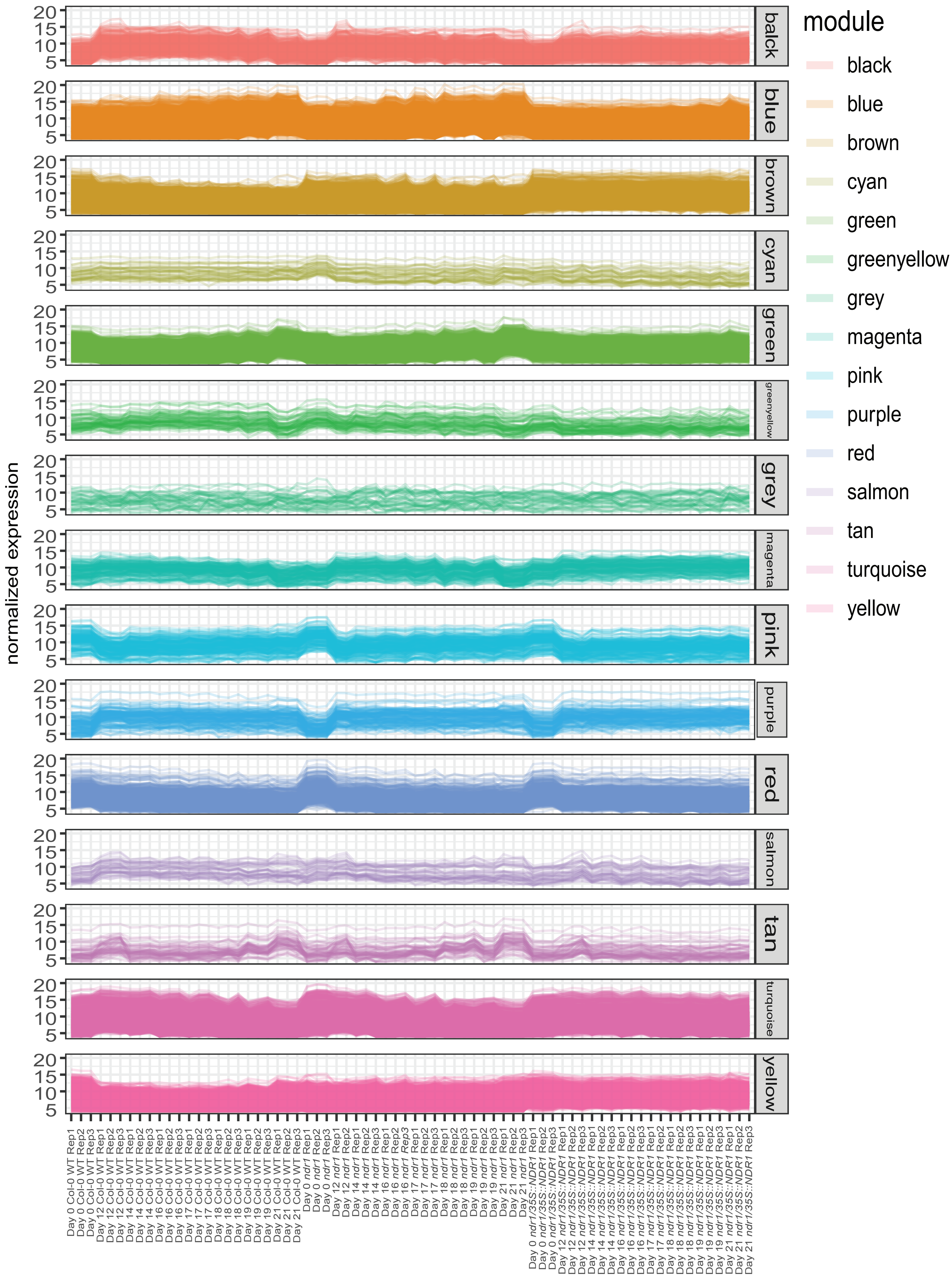
normalized expression
20
15
10
5
balck
blue
brown
cyan
green
greenyellow
grey
magenta
pink
purple
red
salmon
tan
turquoise
yellow
module
black
blue
brown
cyan
green
greenyellow
grey
magenta
pink
purple
red
salmon
tan
turquoise
yellow
Day 0 Col-0 WT Rep1
Day 0 Col-0 WT Rep2
Day 0 Col-0 WT Rep3
Day 12 Col-0 WT Rep1
Day 12 Col-0 WT Rep2
Day 12 Col-0 WT Rep3
Day 14 Col-0 WT Rep1
Day 14 Col-0 WT Rep2
Day 14 Col-0 WT Rep3
Day 16 Col-0 WT Rep1
Day 16 Col-0 WT Rep2
Day 16 Col-0 WT Rep3
Day 17 Col-0 WT Rep1
Day 17 Col-0 WT Rep2
Day 17 Col-0 WT Rep3
Day 18 Col-0 WT Rep1
Day 18 Col-0 WT Rep2
Day 18 Col-0 WT Rep3
Day 19 Col-0 WT Rep1
Day 19 Col-0 WT Rep2
Day 19 Col-0 WT Rep3
Day 21 Col-0 WT Rep1
Day 21 Col-0 WT Rep2
Day 21 Col-0 WT Rep3
Day 0 ndr1 Rep1
Day 0 ndr1 Rep2
Day 0 ndr1 Rep3
Day 12 ndr1 Rep1
Day 12 ndr1 Rep2
Day 12 ndr1 Rep3
Day 14 ndr1 Rep1
Day 14 ndr1 Rep2
Day 14 ndr1 Rep3
Day 16 ndr1 Rep1
Day 16 ndr1 Rep2
Day 16 ndr1 Rep3
Day 17 ndr1 Rep1
Day 17 ndr1 Rep2
Day 17 ndr1 Rep3
Day 18 ndr1 Rep1
Day 18 ndr1 Rep2
Day 18 ndr1 Rep3
Day 19 ndr1 Rep1
Day 19 ndr1 Rep2
Day 19 ndr1 Rep3
Day 21 ndr1 Rep1
Day 21 ndr1 Rep2
Day 21 ndr1 Rep3
Day 0 ndr1/35S::NDR1 Rep1
Day 0 ndr1/35S::NDR1 Rep2
Day 0 ndr1/35S::NDR1 Rep3
Day 12 ndr1/35S::NDR1 Rep1
Day 12 ndr1/35S::NDR1 Rep2
Day 12 ndr1/35S::NDR1 Rep3
Day 14 ndr1/35S::NDR1 Rep1
Day 14 ndr1/35S::NDR1 Rep2
Day 14 ndr1/35S::NDR1 Rep3
Day 16 ndr1/35S::NDR1 Rep1
Day 16 ndr1/35S::NDR1 Rep2
Day 16 ndr1/35S::NDR1 Rep3
Day 17 ndr1/35S::NDR1 Rep1
Day 17 ndr1/35S::NDR1 Rep2
Day 17 ndr1/35S::NDR1 Rep3
Day 18 ndr1/35S::NDR1 Rep1
Day 18 ndr1/35S::NDR1 Rep2
Day 18 ndr1/35S::NDR1 Rep3
Day 19 ndr1/35S::NDR1 Rep1
Day 19 ndr1/35S::NDR1 Rep2
Day 19 ndr1/35S::NDR1 Rep3
Day 21 ndr1/35S::NDR1 Rep1
Day 21 ndr1/35S::NDR1 Rep2
Day 21 ndr1/35S::NDR1 Rep3

**Supplementary Fig. 15 | Modules from coexpression profiles in WT Col-0, *ndr1* and *ndr1/35S::NDR1*.** Samples were collected at 0, 12, 14, 16, 17, 18, 19 and 21 days under well-watered or water withholding conditions.

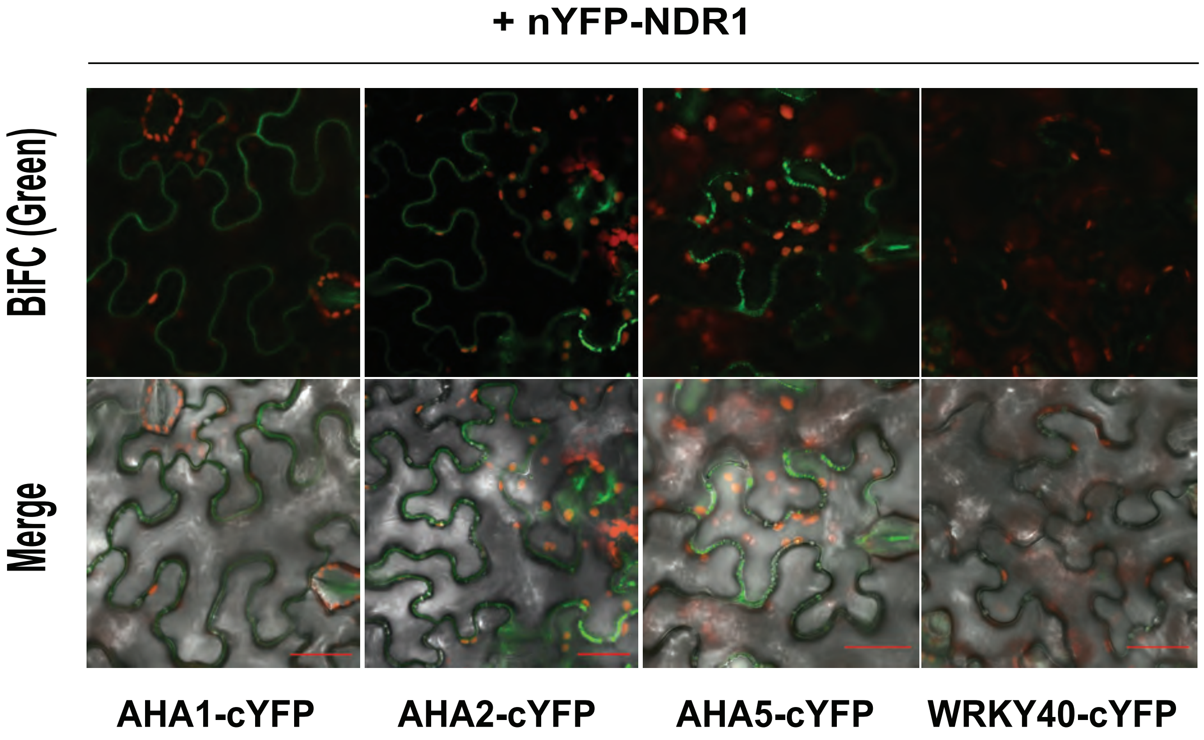


**Supplementary Fig. 16 | BiFC confirmed NDR1 associates with AHA1, AHA2, and AHA5.** Co-expression of n-YFP-NDR1 with $H^+$-ATPase-cYFP proteins in *N. benthamiana* leaves. WRKY40-c-YFP was employed as a negative control.

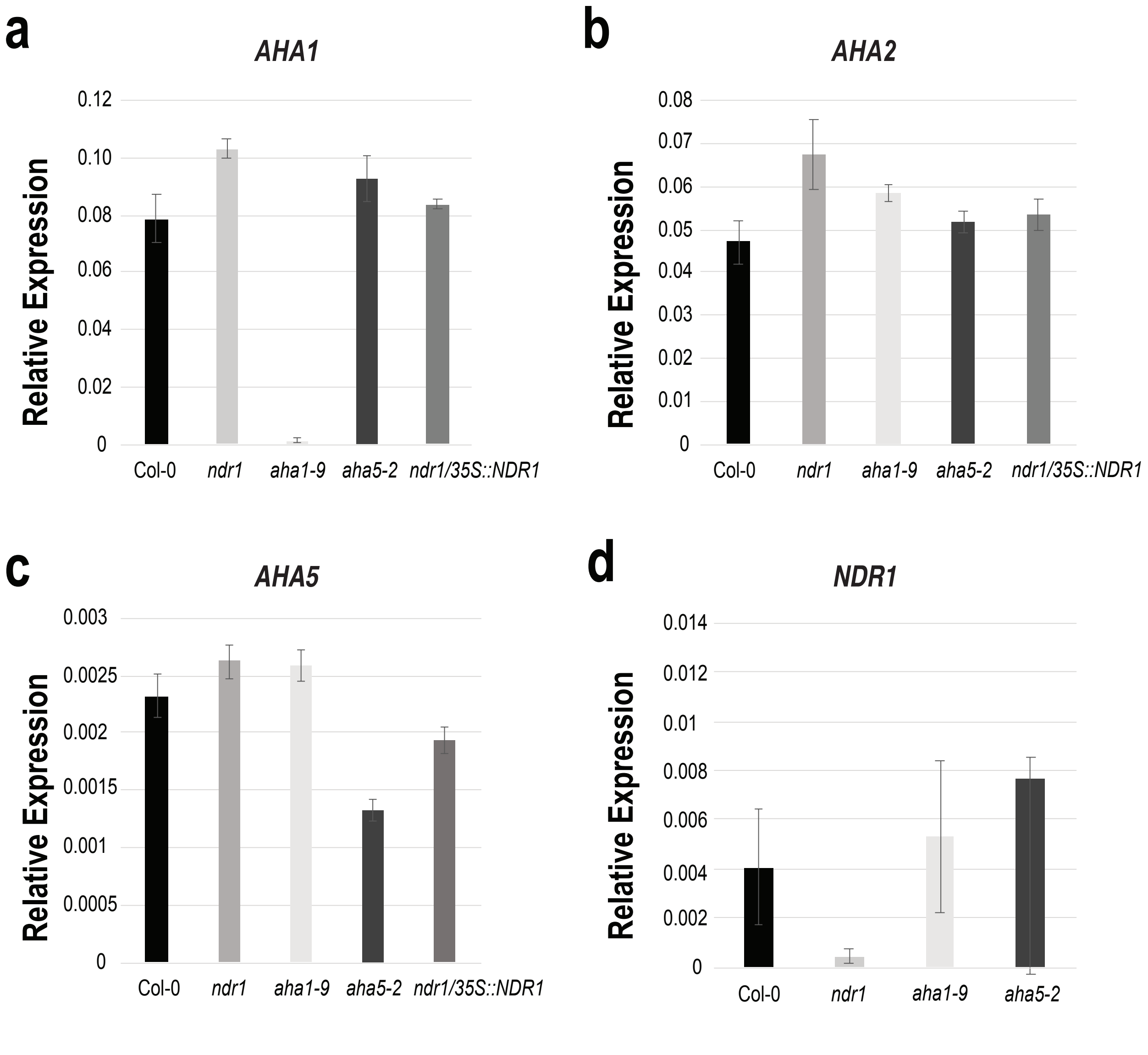


**Supplementary Fig. 17 | mRNA expression levels in 4 Arabidopsis genotypes. a**, *AHA1*, **b,** *AHA2*, **c,** *AHA5*, and **d,** *NDR1*.
